# Rbx1 Cullin-Ring ligase E3 and Ube2m neddylation E2 differentially governs the fitness of Treg cells

**DOI:** 10.1101/2020.07.31.230532

**Authors:** Di Wu, Haomin Li, Mingwei Liu, Jun Qin, Yi Sun

**Affiliations:** Cancer Institute of the Second Affiliated Hospital and Institute of Translational Medicine, Zhejiang University School of Medicine, Hangzhou, Zhejiang, China, 310029; Children’s Hospital, Zhejiang University School of Medicine, Hangzhou, Zhejiang, China, 310003; State Key Laboratory of Proteomics, Beijing Proteome Research Center, National Center for Protein Sciences (Beijing) and Institute of Lifeomics, Beijing, China, 102206

## Abstract

Cullin-RING Ligases (CRLs) are a family of multi-unit E3 ubiquitin ligases with two members of RING family proteins, acting as the catalytic subunit: RING-box 1 (Rbx1) couples with CRLs1-4, whereas RING-box 2 (Rbx2/Sag) couples mainly with CRL5.^1–4^ Activity of CRLs requires neddylation on their Cullin subunit, catalyzed by neddylation enzyme cascades E1, E2 and E3. Ube2m and Ube2f are two neddylation E2s responsible for neddylation of Cullins in CRLs1-4 or CRL5, respectively.^5–6^ Regulatory T cells (Treg cells) are specialized immunosuppressive CD4^+^ T lymphocytes, that play pivotal roles in maintaining immune homeostasis *in vivo*.^7–10^ Whether and how Rbx1-Rbx2/CRLs and Ube2m-Ube2f/neddylation regulate Treg cell homeostasis and function are currently unknown. Here we show that while mice with a Treg-specific deletion of *Rbx2/Sag* showed no obvious phenotype, mice with *Rbx1* deletion in Treg cells developed an early-onset fetal inflammatory disorders and death at day ~25 after birth (~p25), with disrupted homeostasis and functions of Treg cells, indicating Rbx1 as a prominent regulator of Treg cells. Single cell transcriptome assay showed that Rbx1 is essential for the maintenance of the effector subpopulations in Treg cells. The whole genome transcriptome and proteomics analysis revealed that Rbx1 regulates several inflammatory pathways, such as T-cell receptor, IL-17, TNFα, NFκB, chemokine, cytokine-cytokine receptor interaction, as well as energy and purine metabolisms. Accumulation of Acly, Fto and Nfkbib proteins, upon Rbx1 depletion suggests that these are likely the novel substrates of CRLs1-4 in Treg cells. Consistently, while *Ube2f* deletion showed no obvious phenotype, mice with *Ube2m* deletion in Treg cells also suffers from inflammatory disorders, but to a much lesser severity with a 50% of death rate at ~150 days of age. Since Rbx1 is a dual E3 as a component of CRLs1-4 ligase and as a neddylation co-E3, downstream of Ube2m E2 for neddylation activation of CRLs1-4^5^, much severe phenotypes in *Foxp3^cre^*;*Rbx1^fl/fl^* mice suggests *Rbx1* may have additional function independent of neddylation activation in Treg cells. Indeed, unbiased transcriptome comparison between *Rbx1*-deficient and *Ube2m*-deficient Treg cells, revealed that the former had greater as well as unique alteration in the signaling pathways controlling the inflammatory responses. Collectively, our study shows that the Ube2m-Rbx1 axis of the neddylation-CRL is required for the maintenance of homeostasis and functional fitness of Treg cells in the fine control of immune tolerance; with implication that targeting the neddylation/CRLs, such as a small molecule inhibitor pevonedistat^11^, currently in the Phase II clinic trial for anticancer therapy, may have novel application in the treatment of human diseases associated with overactivated Treg cells.

---

A Cullin-RING ligase (CRL) ubiquitin E3 complex consists of 4 subunits: a scaffold Cullin with 8 family members, an adaptor with many members, a substrate receptor with hundreds of members, and a RING catalytic component with two family members, RING-box 1 (Rbx1/Roc1) and RING-box 2 (Rbx2/Roc2/Sag).^12^ Our previous total knockout study showed that *Rbx1* and *Rbx2/Sag* are functionally non-redundant and required for mouse embryogenesis^13–14^. Although mice are viable, T-cell specific *Sag* knockout (via Lck-Cre) significantly decreased T cell activation, proliferation, and T-effector cytokine release^15^, whereas *Sag* knockout in myeloid lineage (via LysM-Cre) increased serum levels of proinflammatory cytokines and enhanced mortality in response to LPS^16^.

To investigate the role of CRLs in regulation of Treg cells *in vivo*, we generated conditional knockout mouse models with inactivation of *Rbx1* and *Rbx2/Sag* individually in Treg cells, driven by *Foxp3^YFP-cre^* (*Foxp3^cre^*)^17,18^. The compound mice were designated as *Foxp3^cre^*; *Rbx1^fl/fl^* or *Foxp3^cre^*;*Sag^fl/fl^* mice, respectively. Compared with *Foxp3^cre^* controls (wild-type/*wt*), *Foxp3^cre^*;*Sag^fl/fl^* mice were viable and healthy without obvious alterations in activation markers of CD4^+^Foxp3^-^ T cells (conventional T cells, or Tcon cells), Tcon cell proliferation rate, and the Treg/CD4^+^ ratio (Extended Data Fig. 1), indicating that Treg specific depletion of Sag does not obviously impair Treg cell function and survival at the steady state.

## Early-onset fatal inflammation in *Foxp3^cre^*;*Rbx1^fl/fl^* mice

Strikingly, deletion of *Rbx1* (confirmed by transcriptome profiling in CD4^+^YFP^+^ Treg cells with 100-fold reduction of *Rbx1* mRNA, Extended data Fig. 2a) in Treg cells was early-onset fatal. Compared with *wt* controls, *Foxp3^cre^*;*Rbx1^fl/fl^* mice showed an altered appearance as early as day 13-15 after birth (p13-15), with collapsed ears, festered skin, and reduced body size (Fig.1a). The mice continued to lose weight dramatically after p15 (Fig.1b) with a 50% or 100% of death rate at p25 or p37, respectively (Fig.1c). Autopsy examination revealed that *Foxp3^cre^*;*Rbx1^fl/fl^* mice had swollen lymph nodes and spleens (Fig. 1d & Extended data Fig. 2b), with lymphocyte infiltration into multiple organs, including the skin, lung, stomach, liver, kidney and colon (Extended data Fig. 2c). The *Foxp3^cre^*;*Rbx1^fl/fl^* mice also had a decreased CD4^+^/CD8^+^ T-cell ratio (Extended data Fig. 2d) and a significantly increased proportion of effector/memory T cells (CD44^hi^CD62L^lo^) among Tcon cells (Fig.1e & Extended data Fig.2e), indicating robust activation of immune cells, a typical phenotype of autoimmune disease. The fatal inflammation observed in *Foxp3^cre^*;*Rbx1^fl/fl^* mice is reminiscent of Treg-deficient mice^10^ or mice with loss-of-function mutations in the *Foxp3* gene^19^, suggesting *Rbx1* is absolutely essential for Treg cells *in vivo*.

**Figure 1.**
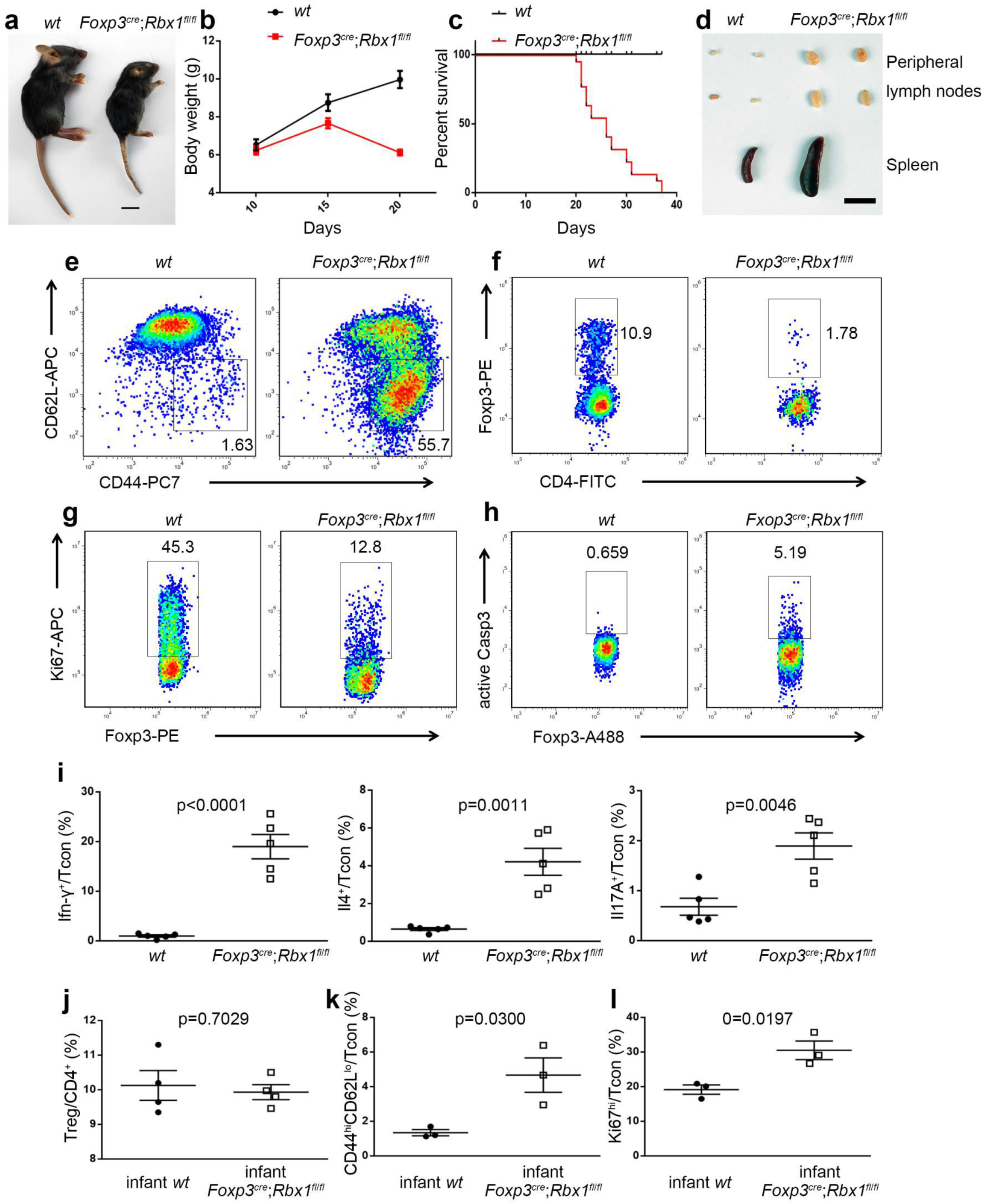
*Rbx1* deletion in Treg cells leads to an early-onset fatal inflammatory disorder. a. Representative images of *wt* and *Foxp3^cre^*;*Rbx1^fl/fl^* mice (p20, scale bar =1cm). b. Gross body weight of *wt* and *Foxp3^cre^*;*Rbx1^fl/fl^* mice (*n* =10). c. Survival curve of *wt* and *Foxp3^cre^*;*Rbx1^fl/fl^* mice (*n* =21; *p* < 0.0001). d. Representative images of the peripheral lymph nodes and spleen from *wt* and *Foxp3^cre^*;*Rbx1^fl/fl^* mice (p22, scale bar = 1 cm). e. Expression of CD44 and CD62L in Tcon cells from peripheral lymph nodes of *wt* and *Foxp3^cre^*;*Rbx1^fl/fl^* mice (p21). f. The proportion of Treg cells among CD4^+^-T cells from peripheral lymph nodes of *wt* and *Foxp3^cre^*;*Rbx1^fl/fl^* mice (p23). g. Expression of Ki67 in Treg cells from peripheral lymph nodes of *wt* and *Foxp3^cre^*;*Rbx1^fl/fl^* mice (p19-23, *n* =5). h. Active Casp3 in Treg cells from peripheral lymph nodes of *wt* and *Foxp3^cre^*;*Rbx1^fl/fl^* mice (p19-23, *n* =4). i. The levels of Ifn-γ, Il-4 and Il-17 in Tcon cells from peripheral lymph nodes in *wt* and *Foxp3^cre^*;*Rbx1^fl/fl^* mice (p19-23, *n* =5). j-l. Impaired suppressive function of Rbx1-deficient Treg cells. j,Treg/CD4^+^ ratios (p8); k, CD44^hi^CD62L^lo^/Tcon ratios (p8); l, Ki67^hi^/Tcon ratios (p8) from peripheral lymph nodes of *wt* and *Foxp3^cre^*;*Rbx1^fl/fl^* infant mice at p8. (*n* =3-4).

## Rbx1 maintains Treg cell homeostasis and suppressive function

In moribund *Foxp3^cre^*;*Rbx1^fl/fl^* mice (p19-23), the Treg cell compartment was reduced (Fig.1f & Extended data Fig.3a), with reduced Ki67 expression (Fig.1g & Extended data Fig.3b) and increased caspase-3 activity (Fig.1h & Extended data Fig.3c), indicating reduced proliferation and increased apoptosis. The levels of Ifn-γ and Il-4 were remarkably increased in Tcon cells by flow cytometry detection of single cell suspension from lymph-nodes (Fig.1i). In addition, the serum concentrations of T_H_1/T_H_2 cytokines (Ifn-γ, Il-2 for T_H_1; Il-4, Il-5, Il6, Il-9, Il-10, and Il-13 for T_H_2) and antibodies (IgA and IgG2a for T_H_1; IgE and IgG1 for T_H_2) were elevated (Extended data Fig.4a&b and Extended data Fig.5a&b), indicating a robust activation of T_H_1/T_H_2-immune responses. For T_H_17 reaction, flow cytometry of single cell suspension from lymph-nodes revealed a 2-fold, but statistically significant increase of Il-17A in Tcon cells, (Fig.1i) and serum concentrations of T_H_17 antibodies (IgG3, IgG2b) were also elevated, although not statistically different (Extended data Fig. 5c). Although the serum concentrations of T_H_17 cytokines (Tnf-α, and GM-Csf) did not change obviously, Il-17A level was reduced, whereas the robust inflammatory storm was evident with significantly increased amount of various cytokines in *Foxp3^cre^*;*Rbx1^fl/fl^* mice (Extended data Fig. 4c&d). Thus, while the T_H_17 reaction was activated modestly, an excessive T_H_1/T_H_2-dominant immune polarization was observed in *Foxp3^cre^*;*Rbx1^fl/fl^* mice.

To determine early alterations, we used the *Foxp3^cre^*;*Rbx1^fl/fl^* mice at p8 stage without apparent inflammatory disorders. While these mice had largely normal proportion of Treg cells among CD4^+^-T cells, the overactivation of T-cells were readily detectable, as evidenced by an increased proportion of effector/memory (CD44^hi^CD62L^lo^) T cells and increased Ki67 levels in Tcon cells (Fig. 1j-l). Thus, Rbx1 is indeed required for the maintenance of suppressive function of Treg cell population.

## Reduction of effector subpopulation of Treg cells upon *Rbx1* deletion

Single-cell RNA sequencing (scRNA-seq) was utilized to unbiasedly dissect the cellular heterogeneity of *wt* and *Rbx1*-deficient CD4^+^YFP^+^ Treg cells from the peripheral lymph nodes of inflammation-free *Foxp3^cre/wt^* and *Foxp3^cre/wt^*;*Rbx1^fl/fl^* mice (Extended data Fig. 6). As revealed in the two-dimensional Uniform Manifold Approximation and Projection (UMAP) of the single-cell transcriptomics data, both *wt* and *Rbx1*-deficient Treg cells could be divided into several subpopulations (Fig.2a & Extended data Fig. 7). Rbx1-depletion increased the subpopulation in Cluster 0, whereas remarkably decreased the subpopulations on Clusters 3 and 4 without affecting other subpopulations obviously (Fig.2b).

**Figure 2.**
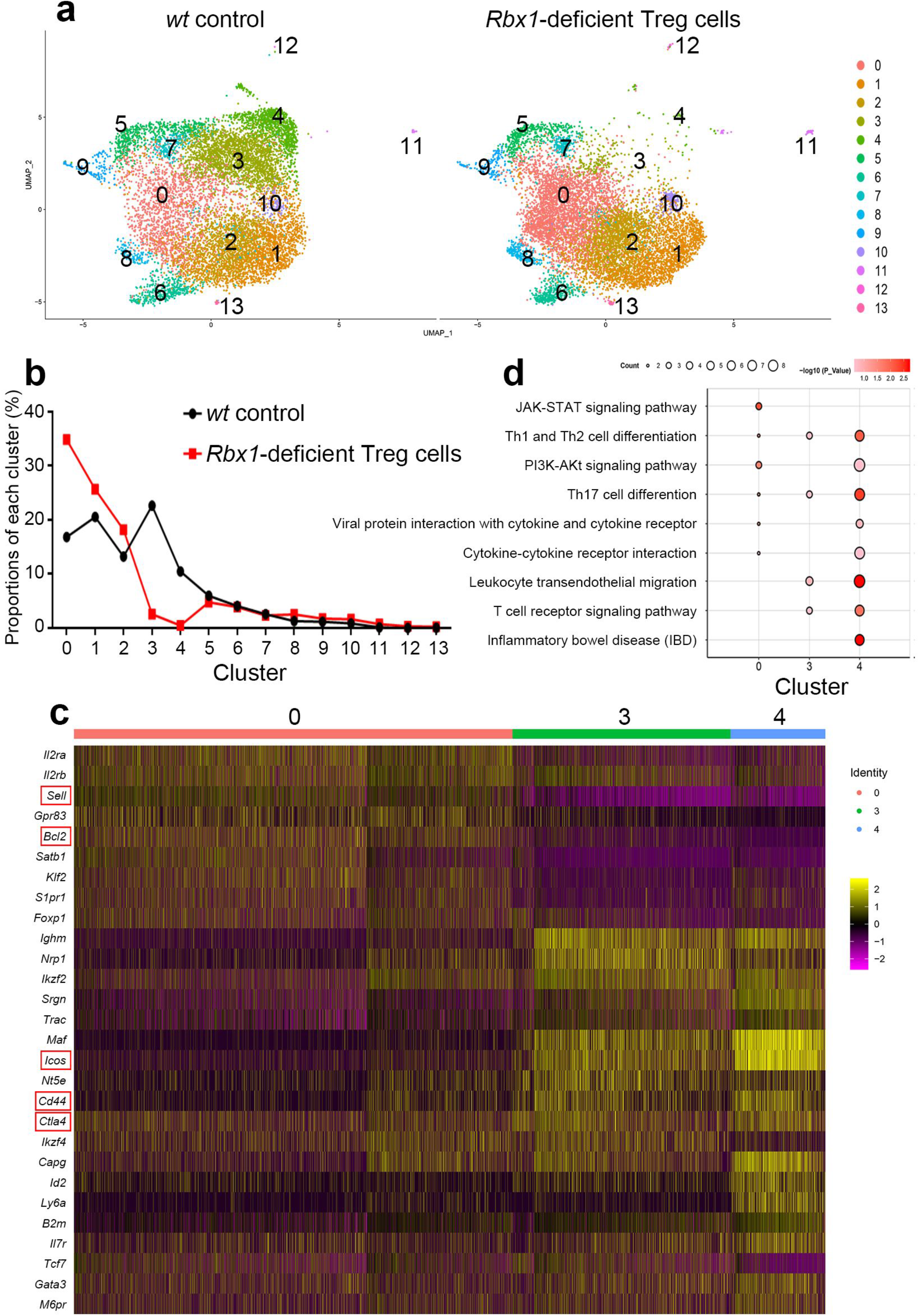
Single-cell transcriptomics of *wt* and *Rbx1*-deficient Treg cells. a. Two-dimensional UMAP visualization of single cell transcriptomics data in CD4^+^YFP^+^ Treg cells from peripheral lymph nodes of *Foxp3^cre/wt^* and *Foxp3^cre/wt^*;*Rbx1^fl/fl^* mice (10 weeks old). b. Proportions of each cluster in CD4^+^YFP^+^ Treg cells from peripheral lymph nodes of *Foxp3^cre/WT^* and *Foxp3^cre/wt^*;*Rbx1^fl/fl^* mice (10 weeks old). c. Heat map of selected marker genes in clusters 0, 3 and 4. Single cells were represented by vertical lines and different colors reflected the relative abundance of indicated genes. d. Selected pathways of clusters 0, 3 and 4 in the KEGG analysis.

The cells in Cluster 0 expressed high levels of genes characteristic of naive Treg cells, including *Sell*^20^ and *Bcl2*; while Treg cell functional molecules, such as *Ctla4*^21^, *Icos*^22^, and *Cd44*^23^, were highly expressed in Cluster 3 and 4 (Fig.2c & Extended data Fig. 8). Thus, Cluster 0 is a quiescent subpopulation and designated as the center population^24^, whereas Clusters 3 and 4 are active Treg cell subpopulations designated as effector populations. The KEGG pathway enrichment analysis revealed that, compared to Cluster 0, T-cell receptor^25^ and other Treg related pathways were activated in Cluster 3 and 4, (Fig. 2d & Extended data Fig. 9). Taken together, Rbx1 is, therefore, essential for the maintenance of effector subpopulations of Treg cells. Lack of effect of Rbx1-deficiency on other subpopulations in Treg cells provides an ideal opportunity of Rbx1 targeting for precise manipulation of Treg cell subpopulations.

## Disordered functional and regulatory network in Rbx1-deficent Treg cells

Transcriptome analysis of Rbx1-deficient Treg cells from inflammation-free *Foxp3^cre/wt^*;*Rbx1^fl/fl^* mice (Extended data Fig. 6) revealed significantly reduced expression of functional genes (Fig. 3a), including *Il10*^17^, *Tgfb*^26^, and *Cst7*^27^ (suppressive cytokines); *Il2r*^28^ and *Entpd1*^29–30^ (regulators of immune cell metabolism); *Ctla4*^22^, *Icos*^23^, *Nrp1*^31^, and *Lag3*^32^ (inhibitor via dendritic cells); and *Fas*^33^ and *Lgals1*^34^ (apoptosis inducer). Impairment of migration-associated surface molecules was also observed in *Rbx1*-deficient Treg cells, including down-regulation of *Cd44, Cxcr3, Cxcr5*, and *Cxcr6*^23^ and up-regulation of *Sell*^21^.

**Figure 3.**
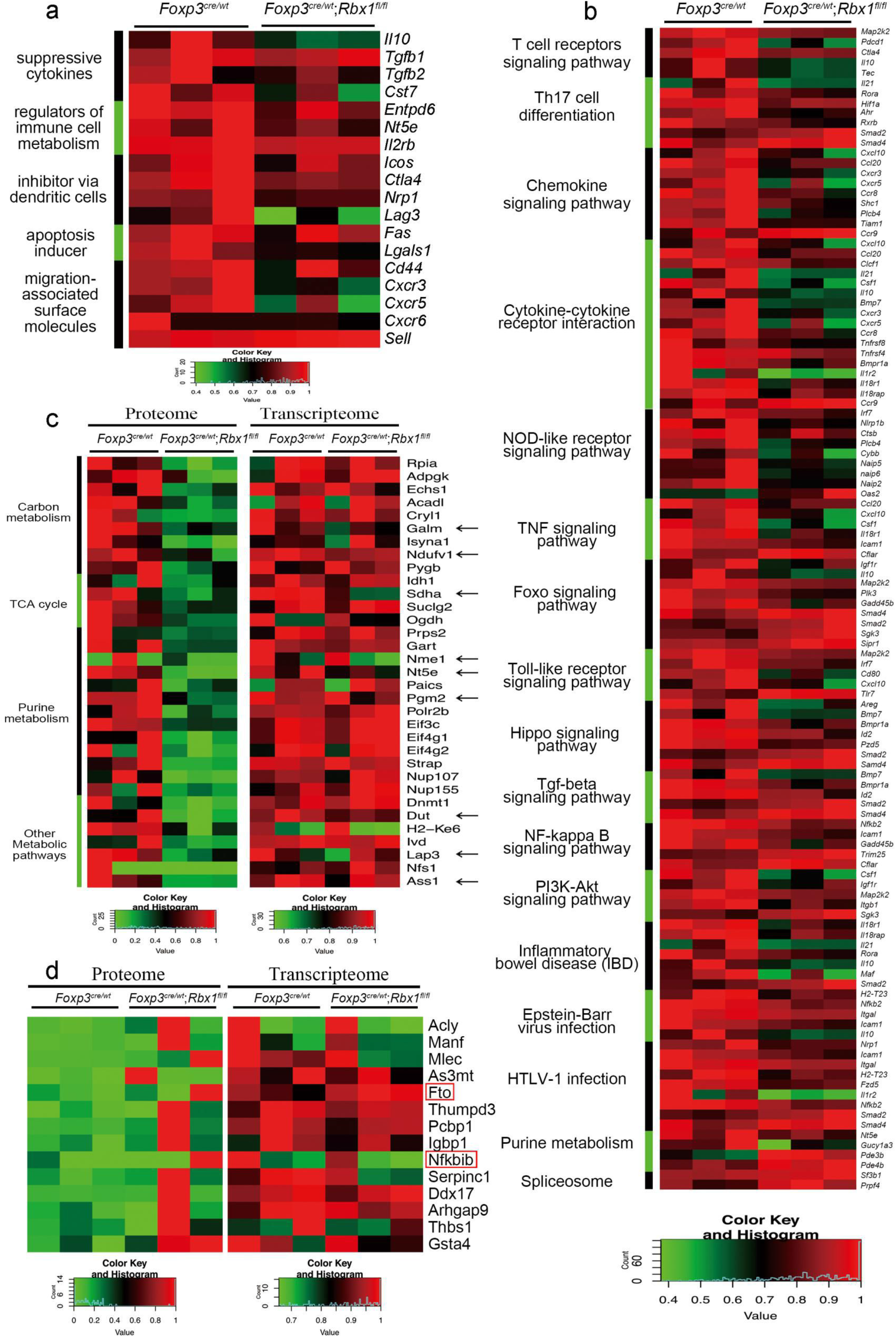
Rbx1-dependent transcriptome and proteome in Treg cells. a. Differentially expressed genes related to Treg function in CD4^+^YFP^+^ Treg cells from *Foxp3^cre/wt^* and *Foxp3^cre/wt^*;*Rbx1^fl/fl^* mice (8-10 weeks old), determined by transcriptional profiling. b. Comparison of expression of genes associated with indicated pathways in CD4^+^YFP^+^ Treg cells derived from *Foxp3^cre/wt^* and *Foxp3^cre/wt^*;*Rbx1^fl/fl^* mice (8-10 weeks old). c. Comparison of the levels of proteins vs. mRNAs in genes associated with indicated pathways in CD4^+^YFP^+^ Treg cells from *Foxp3^cre/wt^* and *Foxp3^cre/wt^*;*Rbx1^fl/fl^* mice (8-10 weeks old). The mRNAs with greater than 1.3-fold changes, which are consistent with protein level changes, were marked by the arrows. d. Comparison of the levels of proteins vs. mRNAs in indicated genes in CD4^+^YFP^+^ Treg cells from *Foxp3^cre/wt^* and *Foxp3^cre/wt^*;*Rbx1^fl/fl^* mice (8-10 weeks old). Those with increased levels of protein but not the mRNA were considered as candidates of the direct degradation substrates of Rbx1 E3 ligase in Treg cells.

The KEGG pathway enrichment analysis revealed that Rbx1 depletion caused downregulation of transcriptome involved in many signaling pathways, including T-cell receptor, Th17 cell differentiation, chemokine, cytokine-cytokine receptor interaction, NOD-like receptor, TNF, FoxO, Toll-like receptor, Hippo, TGFβ, NF-κB, PI3K-AKT, and purine metabolism among few other virus associated pathways (Fig. 3b & Extended data Fig. 10). The results strongly suggest Rbx1 regulation of numerous genes controlling the inflammation, immunological responsiveness, proliferation/survival, and metabolism.

## Specific spectrum of Rbx1 substrates in Treg cells

Considering Rbx1 is an E3 ubiquitin ligase that directly controls the abundance of substrate proteins, we utilized the mass spectrometry-based proteomics to profile the difference in protein amounts between Wt and Rbx1-deficient CD4^+^YFP^+^ Treg cells (three independent sets of combined samples) from inflammation-free *Foxp3^cre/wt^* and *Foxp3^cre/wt^*;*Rbx1^fl/fl^* mice (Extended data Fig. 6). Among a total of 3236 proteins detected in any one of 6 paired samples, we identified 43 increased and 78 decreased proteins with a 3-fold cut-off, and the numbers increased to 70 and 195, respectively, with a 2-fold cut-off in Rbx1-deficient CD4^+^YFP^+^ Treg cells, as compared to the wild-type. The KEGG pathway enrichment analysis of 2-fold cut-off revealed that Rbx1 depletion caused accumulation of few proteins associated with the splicesome or complement and coagulation cascade (Extended data Fig. 11), but significant reduction of proteins controlling TCA cycle, carbon and purine metabolisms, along with other metabolic pathways (Fig. 3c), suggesting a requirement of Rbx1 in regulation of energy and nucleotide metabolisms of Treg cells. We noticed that less than half of protein changes seen in the proteomic analysis matched with the mRNA changes by the transcriptomic analysis (Fig. 3c & Extended data Fig. 11), which is slightly lower than the general correlation of 50% between proteome and transcripteome^35^.

To evaluate the direct substrates of Rbx1 ubiquitin E3 ligase in Treg cells, the proteomic and transcriptomic data were combined for analysis. The candidates are those with increased levels of protein but not the mRNA in Rbx1-deficient Treg cells, which include Acly, Manf, Mlec, As3mt, Fto, and Nfkbib, among the others (Fig. 3d). Acly^36^ and Nfkbib/IκB-β^37^ were previously reported as Rbx1 substrates, whereas the others are likely the novel substrates. Interestingly, many common substrates of Rbx1/CRLs1-4^38–39^ were not detected in Rbx1 deficient Treg cells, few detected ones, like Cdkn1b/p27^13^ and Foxo1^40^, were not accumulated (Extended data Fig. 12), indicating that a subset of Rbx1 substrates are selectively expressed, and specifically subjected to Rbx1 modulation in Treg cells.

Among the putative substrates of Rbx1-CRLs1-4 functioning uniquely in Treg cells, Nfkbib/IκB-β is the inhibitor of NF-κB pathway, whereas Fto is the demethylation enzyme of m^6^A mRNA methylation. NF-κB pathway^18,41^ and m^6^A mRNA methylation^42^ are known to regulate Treg cell functions. Their accumulation upon Rbx1 depletion is likely contributing to observed malfunctions of Treg cells.

## Rbx1 acts in Treg cells via Ube2m-dependent and-independent mechanisms

It has been well-established that the E3 ligase activity of the CRLs requires neddylation of the scaffold protein Cullin^6^. Specifically, neddylation E2 Ube2m couples with Rbx1 (neddylation co-E3) selectively promotes neddylation on Cullins1-4, which are associated with Rbx1 to constitute active CRLs1-4 ubiquitin E3; whereas neddylation E2 Ube2f couples with Sag/Rbx2 E3 to promote neddylation of Cullin5, which is associated with Rbx2 to constitute active CRL5^5^. We, therefore, asked whether the pivotal role of Rbx1 in Treg cells is dependent on Ube2m, whereas Ube2f is non-essential for Treg cells, given the non-essentiality of Sag/Rbx2. Indeed, *Foxp3^cre^*;*Ube2f ^fl/fl^* mice are viable and develop normally without any abnormality (Extended data Fig. 13), indicating that the Ube2f-Rbx2-CRL5 axis is not essential for the functionality and survival of Treg cells at the steady status.

Consistently, like *Foxp3^cre^*;*Rbx1^fl/fl^* mice, *Foxp3^cre^*;*Ube2m^fl/fl^* mice with *Ube2m* deletion in Treg cells (again confirmed by transcriptome profiling, Extended data Fig. 14a), displayed abnormal appearances such as collapsed ears and festered skin, but at the much later stage (Fig. 4a). The growth of mice was remarkably retarded (Fig. 4b), and about 50% of the mice dies at age of 150 days and a 100% death rate within one year (Fig. 4c). The swollen spleens and lymph nodes were apparent (Fig. 4d & Extended data Fig.14b). *Foxp3^cre^*; *Ube2m^fl/fl^* mice also showed obvious T_H_1/T_H_2-polarized immune over-activation but in a much lower levels of inflammation than that in *Foxp3^cre^*;*Rbx1^fl/fl^* mice (Fig.4e&f, Extended data Fig.14-16), but had increased Treg/CD4^+^-T ratio, and lack of Ki67-decrease and active-Casp3-increase in Treg cells (Extended data Fig.17). These observations demonstrate the similarities and differences between the functions of Rbx1 and Ube2m in Treg cells.

**Figure 4.**
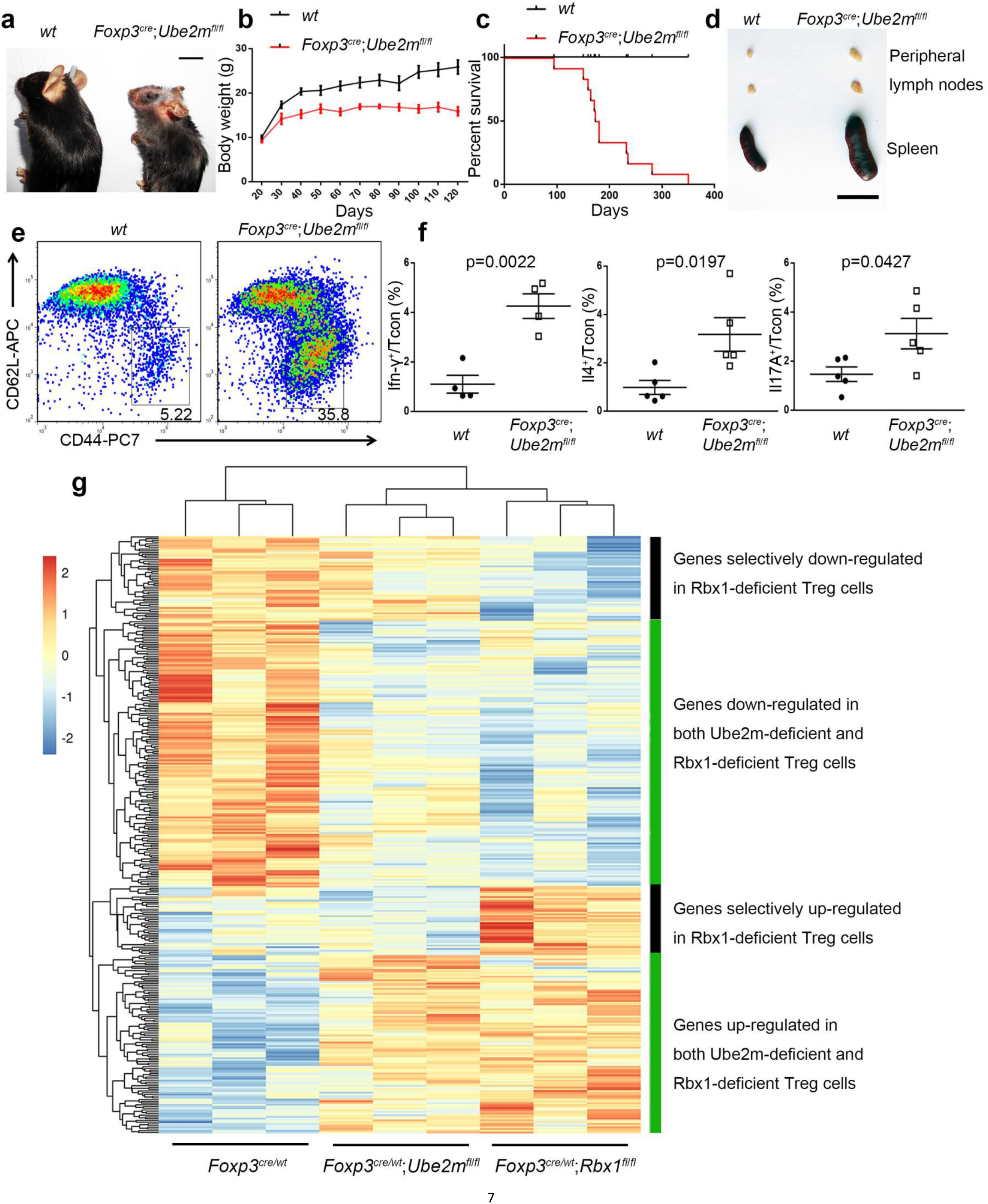
Ube2m-dependent and -independent mechanisms of Rbx1 in Treg cells. a. Representative images of *wt* and *Foxp3^cre^*;*Ube2m^fl/fl^* mice (16 weeks old, scale bar = 1cm). b. Gross body weight of *wt* and *Foxp3^cre^*;*Ube2m^fl/fl^* mice (*n* =20). c. Survival curve of *wt* and *Foxp3^cre^*;*Ube2m^fl/fl^* mice (*n* =12; *p* < 0.0001). e. Representative images of the peripheral lymph nodes and spleen from *wt* and *Foxp3^cre^*;*Ube2m^fl/fl^* mice (16 weeks old, scale bar = 1cm). e. Expression of CD44 and CD62L in Tcon cells from peripheral lymph nodes of *wt* and *Foxp3^cre^*; *Ube2m^fl/fl^* mice (16 weeks old). f. The levels of Ifn-γ, Il-4 and Il-17 in Tcon cells from peripheral lymph nodes of *wt* and *Foxp3^cre^*;*Ube2m^fl/fl^* mice (16 weeks old, *n* =4) g. Unbiased cluster analysis of the transcriptional programs revealed 4 categories of genes differentially expression in *Rbx1*- and *Ube2m*-deficient Treg cells, as compared to the *wt* control Treg cells.

The genome-wide transcriptome analyses of *Ube2m*-deficient Treg cells, as compared to *Rbx1*-deficient Treg cells, revealed the similar changes in the Treg functional genes (Extended data Fig.18), controlling multiple signaling pathways involved in inflammation responses (Extended data Fig.19), but generally to a lesser extent. An unbiased cluster analysis of the transcriptomes between *Rbx1*-deficient and *Ube2m*-deficient Treg cells revealed a good overall correction, but two sets of data did have unique changes in several groups of genes specifically seen in *Rbx1*-deficient Treg cells (Fig.4g and Extended data Fig.20&21), suggesting the dependent and independent mechanisms between Rbx1 and Ube2m in regulation of Treg cell functions. Taken together, both the phenotype comparison and unbiased cluster analyses of the transcriptome support the notion that Rbx1 acts via Ube2m-dependent and -independent mechanisms in Treg cells.

## Discussion

Many preclinical studies have demonstrated that overactivation of protein neddylation and CRL E3s are common in human cancers, which is actively involved in promoting tumorigenesis.^12^ Pevonedistat (also known as MLN4924), a small molecule inhibitor of neddylation E1, that blocks the entire protein neddylation and inhibites all CRL E3s, is currently in several phase II clinical trials for anti-cancer application.^39^ We showed here that inactivation of neddylation by depletion of Ube2m E2 or inactivation of CRLs1-4 by depletion of Rbx1 in Treg cells dramatically inhibits functions and survival of Treg cells, leading to development of the autoimmune disorders with overactivated T toxic cells and premature death. The Treg Rbx1-deficient mice have much severer phenotype than the Ube2m-deficient mice suggest Rbx1/CRLs1-4 may have additional function beyond neddylation activation. Lack of phenotype by *Rbx2*/*Sag* or *Ube2f* deletion in Treg cells excludes possible involvement of CRL5 in Treg regulation.

While both Rbx1 and Ube2m controls the expression of genes regulating several inflammation-related pathways (Extended data Fig.19), the downregulation of *Egr2*^43^, *Ccr8*^44^ and *Tigit*^45^ appear to be relatively unique to Rbx1-deficient Treg cells (Extended data Fig. 20). Altered expression of these genes, which has been shown to regulate Treg cell function, would likely be responsible for the more severe phenotypes seen in *Foxp3^cre^*;*Rbx1^fl/fl^* mice. The KEGG pathway analysis of a group of genes uniquely responded to *Rbx1*-deficiency revealed Rbx1 regulation of several biosynthesis processes and cell-cell adherent junctions, among few others, (Extended data Fig.21) which is an interesting subject for future investigation. More important, Rbx1 was found to control several metabolic pathways, particularly purine pathways shown in both proteomics and transcriptomics analyses (Fig. 3c), suggesting reduced operation of these pathways upon Rbx1 depletion may also contribute to the loss of Treg function and eventually death of animals.

In summary, our study revealed that Rbx1 is essential for the functionality and survival of Treg cells by controlling multiple inflammatory and proliferation/survival signaling, as well as metabolic pathways. Our study also provided a piece of proof-of-concept evidence that the neddylation inhibitor pevonedistat or selective targeting Treg-RBX1, may have novel application in the treatment of human diseases associated with overactivation of Treg cells by inactivating of neddylation/CRLs.

## Acknowledgments

We would like to thank Prof. Xiaoming Feng for providing *Foxp3^YFP-cre^* mice. We also thank the Core Facilities, Zhejiang University School of Medicine for the technical support. This project was supported by the National Key R&D Program of China (2016YFA0501800 to Y. S.); National Science Foundation of China (81801567 to D. W. and 81572718 to Y. S.).

## Author contributions

Di Wu designed and preformed experiments, analyzed data, and wrote the manuscript. Haomin Li analyzed proteome and transcriptome data. Mingwei Liu performed the mass spectrum experiment and analyzed the data. Jun Qin designed and supervised the mass spectrum experiment and analyzed data. Yi Sun designed experiments, analyzed data, wrote the manuscript, and oversaw the project.

## METHODS

### Data reporting

No statistical methods were used to predetermine sample size. The experiments were not randomized and the investigators were not blinded to allocation during experiments and outcome assessment.

### Mice

The *Foxp3^YFP-cre^* mice (Jax No. 016959) are gifts from Prof. Xiaoming Feng (Peking Union Medical College, China).

The *Rbx1^fl/fl^* mice were generated on the C57BL/6 background via CRISPR/Cas9 technology with exons 2-4 floxed (performed by Nanjing Biomedical Research Institute of Nanjing University). Briefly, the sgRNA (5’-A CCACTAAGTGGATAAT CAC-3’ and 5’-GGAGTCAGAAATGATTGGTC-3’) direct Cas9 endonuclease cleavage in intron1-2 and intron 4-5 to insert LoxP sites by homologous recombination. The PCR genotyping primers 5’-GATTCTACTTTGCTTGCAGTGC TC-3’ and 5’-TCTGCATAAGCACGGGCTCTC-3’ generated DNA fragment of 266bp and 196bp bands, respectively, for mutant and wild type.

The *Ube2m^fl/fl^* mice were generated on the C57BL/6 background via CRISPR/Cas9 technology with exons 2-4 floxed (performed by Nanjing Biomedical Research Institute of Nanjing University). Briefly, the sgRNA (5’-TCAATTTAAGCTACCATA C-3’ and 5’-AGTGAAGATACGGGACA-3’) direct Cas9 endonuclease cleavage in intron1-2 and intron 4-5 to insert LoxP sites by homologous recombination. The PCR genotyping primers 5’-CAAGACCTGCCTTCCAGGTATC-3’ and 5’-CCCTGCTAA TACTGAACAGG-3’ generated DNA fragment of 317bp and 223bp for mutant and wildtype, respectively,

The *Ube2f^fl/fl^* mice were generated from an ES clone (HEPD0820_4_G09), purchased from EuMMCR (European Mouse Mutant Cell Repository) (MGI:1915171) with exon 5 floxed. For LoxP sites flanked allele, the PCR genotyping primers are 5’-GCGAGCTCAGACCATAACTTCG-3’ and 5’-CCAGGGTGGAAAATTTCAGT TT-3’; and for wild type allele, the genotyping primers are 5’-CCCTGGAATTTCGG TATTATA-3’ and 5’-CCAGGGTGGAAAATTTCAGTTT-3’, respectively.

The *Sag^fl/fl^* mice were generated and characterized as described^46^.

All mice were maintained in SPF conditions. All animal experiments were approved by the Animal Ethics Committee of Zhejiang University; animal care was provided in accordance with the principles and procedures by the regulatory standards at Zhejiang University Laboratory Animal Center.

### Flow cytometry

For analysis of surface markers, cells were washed in PBS containing 2% (wt/vol) FBS, and stained with indicated Abs according to the manufacturer’s instructions (BD Pharmingen, 562574), followed by flow cytometry analysis. For intracellular cytokine staining, cells were stimulated for 5 h with leukocyte activation cocktail of GolgiPlug (BD Pharmingen, 550583). Antibodies were all from eBioscience, unless otherwise noted: anti-CD4 (GK1.5), anti-CD8α (53-6.7, BD Pharmingen), anti-CD44 (IM7, BD Pharmingen), anti-CD62L (MEL-14), anti-Foxp3 (FJK-16s), anti-Ki67 (SolA15), anti-active caspase 3 (C92-605, BD Pharmingen), anti-Ifn-γ (XMG1.2), anti-Il4 (11B11), anti-Il17A (eBio17B7). Flow cytometry data were acquired on CytoFLEX LX or CytoFLEX S (Beckman) and analyzed using Flowjo software (Tree Star).

### Cytokines assay

The serum was drawn from mice of indicated genotypes, diluted 4-fold by PBS, and then subjected to cytokine/chemokine profiling using Mouse Cytokine Grp I (Bio-Plex Pro, M60009RDPD), measured on Luminex 200 system.

### Antibodies assay

The immunoglobulin subclasses in mouse serum were measured using Mouse Immunoglobulin Isotyping Kit (BD Cytometric Bead Array, 550026).

### Cell sorting

Peripheral lymph nodes and/or spleens were harvested from mice with indicated genotypes, and grinded into single cells, followed by treatment with Dynabeads Untouched Mouse CD4 Cells Kit (Invitrogen, 11415D) to isolate the CD4^+^ T cells. The FACS sorting (SONY Cell Sorter, SH800S) was used to isolate the CD4^+^YFP^+^ Treg cells with purities > 99%.

### Single-cell RNA sequencing

CD4^+^YFP^+^ Treg cells were isolated from the peripheral lymph nodes of mice with indicated genotypes as above. To generate enough materials, the lymph nodes from 3 *Foxp3^cre/wt^* or 6 *Foxp3^cre/wt^*;*Rbx1^fl/fl^* mice at age of 10 weeks were pooled and subjected to single-cell RNA sequencing by Sinotech Genimics Co., Ltd. (Shanghai, China). The single-cell capture was achieved by random distribution of a single-cell suspension across > 200,000 microwells through a limited dilution approach, and the *wt* and *Rbx1*-deficient Treg cells were distributed on two individual microwell plates. Beads with oligonucleotide barcodes were added to saturation so that a bead was paired with a cell in a microwell. Cell-lysis buffer was added to facilitate poly-adenylated RNA molecules hybridized to the beads. Beads were collected into a single tube for reverse transcription. Upon cDNA synthesis, each cDNA molecule was tagged on the 5’ end (that is, the 3’ end of a mRNA transcript) with a molecular index and cell label to indicate its cell of origin. Whole transcriptome libraries were prepared using the BD Resolve system for single-cell whole-transcriptome amplification workflow (BD Genomics) to capture transcriptomic information of the sorted single cells. Briefly, the second strand cDNA was synthesized, followed by ligation of the adaptor for universal amplification. Eighteen cycles of PCR were used to amplify the adaptor-ligated cDNA products. Sequencing libraries were prepared using random priming PCR of the whole-transcriptome amplification products to enrich the 3’ end of the transcripts linked with the cell labels and molecular indices. Sequencing libraries were quantified using a High Sensitivity DNA chip (Agilent) on a Bioanalyzer 2100 and the Qubit High Sensitivity DNA assay (Thermo Fisher Scientific). Approximate 1.5pM of the library for each sample was loaded onto a NextSeq 500 system and sequenced using High Output sequencing kits (75× 2bp) (Illumina). After discarding the cells with high mitochondrial gene, 10,048 and 9,736 cells were identified for *wt* and *Rbx1*-deficient Treg cells, with an average number of reads of 23,321.27 vs. 22202.55, and 1,080.16 vs. 870.28 of genes detected per cell, respectively.

### Transcriptome profiling

CD4^+^YFP^+^ Treg cells were isolated from the peripheral lymph nodes and spleens of mice (8-12 weeks old) with indicated genotypes. To generate enough materials, the pooled tissues from 2-3 mice (*Foxp3^cre/wt^*), 8-11 (*Foxp3^cre/wt^*;*Rbx1^fl/fl^*) or 3-4 (*Foxp3^cre/wt^*;*Ube2m^fl/fl^*) were used. RNA samples were prepared with the miRNeasy Mini Kit (Qiagen, 21704), then reverse transcribed, amplified, labeled (Affymetrix GeneChip pico kit, 703308), and hybridized to Clariom S Arrays, mouse (Affymetrix, 902931), performed by Cnkingbio Biotech Co., Ltd. (Beijing, China). Microarray data sets were analyzed with Applied Biosystems Expression Console Software 1.4. Three independent experiments were performed.

### Proteomics profiling

CD4^+^YFP^+^ Treg cells were isolated from the peripheral lymph nodes and spleens of mice (>8 weeks of age) with indicated genotypes. To secure enough samples, tissues from 5-6 of *Foxp3^cre/wt^* or 21-24 of *Foxp3^cre/wt^*;*Rbx1^fl/fl^* mice were pooled, washed with PBS buffer, and stored at −80°C. Three independent collected pooled samples were subjected to Mass-Spectrometry.

Cells were resuspended in 5μl 1% (w/v) sodium deoxycholate, 10mM TCEP, 40mM 2-chloroacetamide (CAA), 100mM Tris (pH 8.5), and subsequently lysed by 5min boiling at 95°C and sonication (Bioruptor, model UCD-200, Diagenode) for 15min. Cell debris were pelleted by centrifugation at 13,200r.p.m. for 5min and the clarified lysate was transferred into a new vial. The lysate was diluted 1:10 for LysC-trypsin digestion (0.4μg of each enzyme in double distilled water), and the digestion was performed overnight at 37°C. The digest was acidified with 50μl 2% TFA and sodium deoxycholate was extracted using 50μl ethyl acetate and vigorous shaking. The organic phase was removed after centrifugation at 13,200r.p.m. for 5min. Finally, the peptides were desalted on C18 StageTips.

Samples were analyzed on Orbitrap Fusion Lumos mass spectrometers (Thermo Fisher Scientific, Rockford, IL, USA) coupled with an Easy-nLC 1000 nanoflow LC system (Thermo Fisher Scientific). Dried peptide samples were re-dissolved in Solvent A (0.1% formic acid in water) and loaded to a trap column (100μm × 2cm, home-made; particle size, 3μm; pore size, 120Å; Dr. Maisch, Ammerbuch, Germany) with a max pressure of 280 bar using Solvent A, then separated on a home-made 150μm×30cm silica microcolumn (particle size, 1.9μm; pore size, 120Å; Dr. Maisch, Ammerbuch, Germany) with a gradient of 5-35% mobile phase B (acetonitrile and 0.1% formic acid) at a flow rate of 600 nl/min for 141 min then up to 95% in 1 minute and eluted for 9min (the column temperature was maintained at 60°C). The MS analysis were performed in data-dependent acquisition mode. Automatic gain control (AGC) targets were 5×10^5^ ions with a max injection time of 50 ms for full scans and 5×10^3^ with 35 ms for MS/MS scans. The most intense ions selected under top-speed mode were isolated in Quadrupole with a 1.6m/z window and fragmented by higher energy collisional dissociation (HCD) with normalized collision energy of 32% and dynamic exclusion time was set as 25s. Data were acquired using the Xcalibur software (Thermo Fisher Scientific).

DAVID was used for pathway enrichment analysis of proteins with change fold > 2/3 or < 0.5/0.33. The enriched pathways and their contained proteins with 2-fold change were shown in heat map in which each quantitative value was normalized based its maximum value.

### Statistical analysis

The *p* values were calculated by Mann–Whitney test, two-tailed unpaired Student’s t-test, one-way ANOVA or two-way ANOVA as indicated using GraphPad Prism, unless otherwise noted. Statistical analysis of mouse survival and respective *p* values were determined using the log-rank test. *p* < 0.05 was considered as significant. All error bars represent the s.e.m. from three independent experiments.

### Data availability

The scRNA-seq and microarray data that support the findings of this study will be deposited in the Gene Expression Omnibus.

