## Extended figures for "Rbx1 Cullin-Ring ligase E3 and Ube2m neddylation E2 differentially governs the fitness of Treg cells"

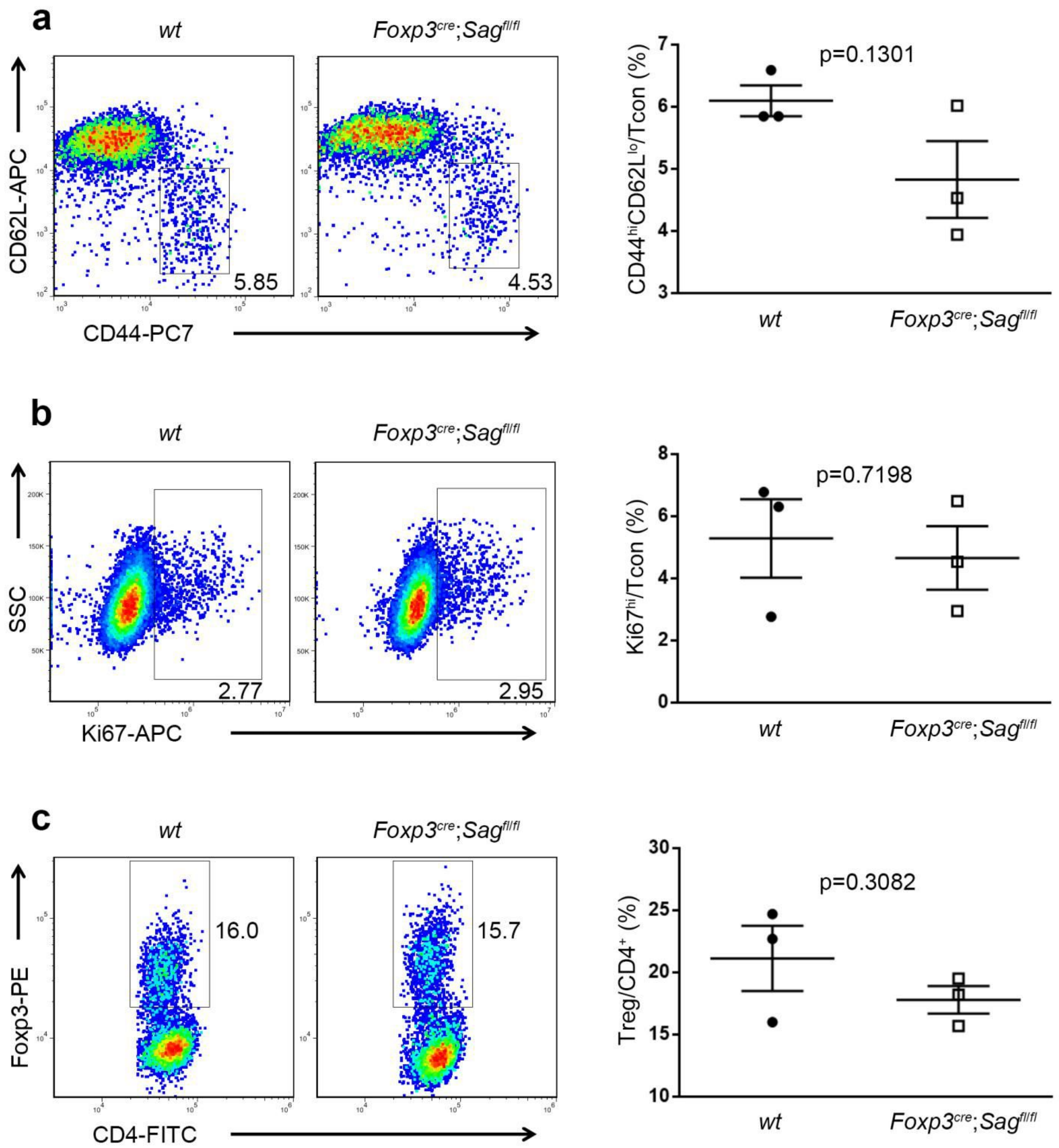


**Extended Data Figure 1. Deficiency of *Rbx2/Sag* does not obviously impair Treg cell fitness at the steady status**

a. Expression of CD44 and CD62L in Tcon cells from peripheral lymph nodes of *wt* and *Foxp3*$^{cre}$;*Sag*$^{fl/fl}$ mice (10 weeks old, *n*=3).

b. Expression of Ki67 in Tcon cells from peripheral lymph nodes of *wt* and *Foxp3*$^{cre}$;*Sag*$^{fl/fl}$ mice (10 weeks old, *n*=3).

c. The proportion of Treg cells among CD4$^{+}$-T cells from peripheral lymph nodes of *wt* and *Foxp3*$^{cre}$;*Sag*$^{fl/fl}$ mice (10 weeks old, *n*=3).

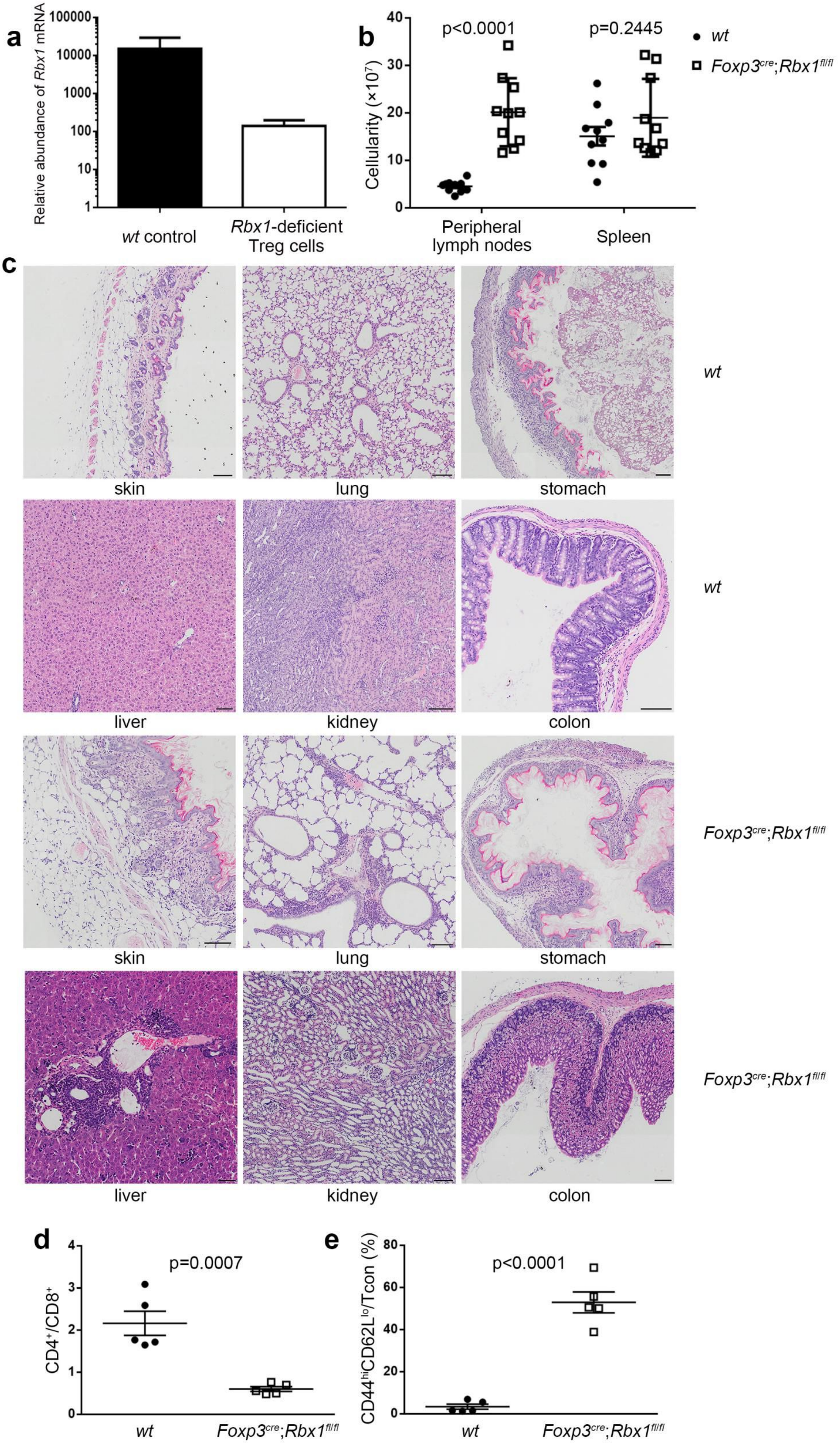
a
Relative abundance of Rbx1 mRNA
100000
10000
1000
100
10
1
wt control
Rbx1-deficient Treg cells
b
Cellularity (×10⁷)
40
30
20
10
0
p<0.0001
p=0.2445
wt
Foxp3cre;Rbx1fl/fl
Peripheral lymph nodes
Spleen
c
skin
lung
stomach
wt
liver
kidney
colon
wt
skin
lung
stomach
Foxp3cre;Rbx1fl/fl
liver
kidney
colon
Foxp3cre;Rbx1fl/fl
d
CD4+/CD8+
4
3
2
1
0
p=0.0007
wt
Foxp3cre;Rbx1fl/fl
e
CD44hiCD62Llo/Tcon (%)
80
60
40
20
0
p<0.0001
wt
Foxp3cre;Rbx1fl/fl

**Extended Data Figure 2. Disastrous auto-immune disorders in *Foxp3*$^{cre}$;*Rbx1*$^{fl/fl}$ mice**

a. Relative *Rbx1* mRNA abundance in *wt* and *Rbx1*-deficient Treg cells revealed by transcriptome profiling.

b. Total cell numbers in peripheral lymph nodes and spleen from *wt* and *Foxp3*$^{cre}$;*Rbx1*$^{fl/fl}$ mice (p19-23, *n*=10).

c. H&E staining of the skin, lung, stomach, liver, kidney, and colon from *wt* and *Foxp3*$^{cre}$;*Rbx1*$^{fl/fl}$ mice (p19-20, scale bar = 50μm in liver, or 100μm in other organs).

d. $CD4^+/CD8^+$ ratios in peripheral lymph nodes from *wt* and *Foxp3*$^{cre}$;*Rbx1*$^{fl/fl}$ mice (p19-23, *n* =5).

e. $CD44^{hi}CD62L^{lo}$/Tcon ratios in peripheral lymph nodes from *wt* and *Foxp3*$^{cre}$;*Rbx1*$^{fl/fl}$ mice (p19-23,*n* =5).

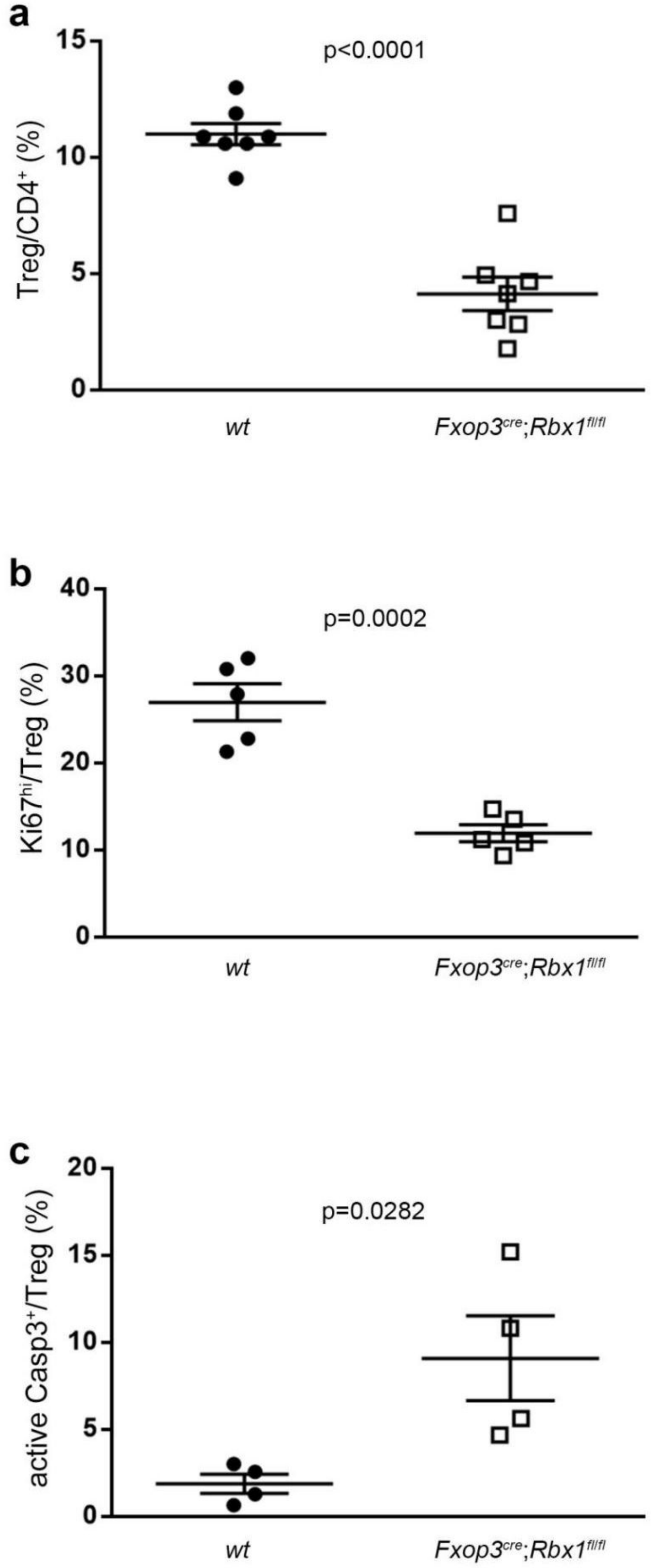


**Extended Data Figure 3. Reduced proliferation and elevated apoptosis in Rbx1-deficient Treg cells**

a. Treg/CD4$^{+}$ ratios in peripheral lymph nodes from *wt* and *Foxp3*$^{cre}$;*Rbx1*$^{fl/fl}$ mice (p19-23, *n* =7).

b. Ki67$^{hi}$/Treg ratios in peripheral lymph nodes from *wt* and *Foxp3*$^{cre}$;*Rbx1*$^{fl/fl}$ mice (p19-23, *n* =5).

c. Active-Casp3$^{+}$/Treg ratios in peripheral lymph nodes from *wt* and *Foxp3*$^{cre}$;*Rbx1*$^{fl/fl}$ mice (p19-23, *n* =4).

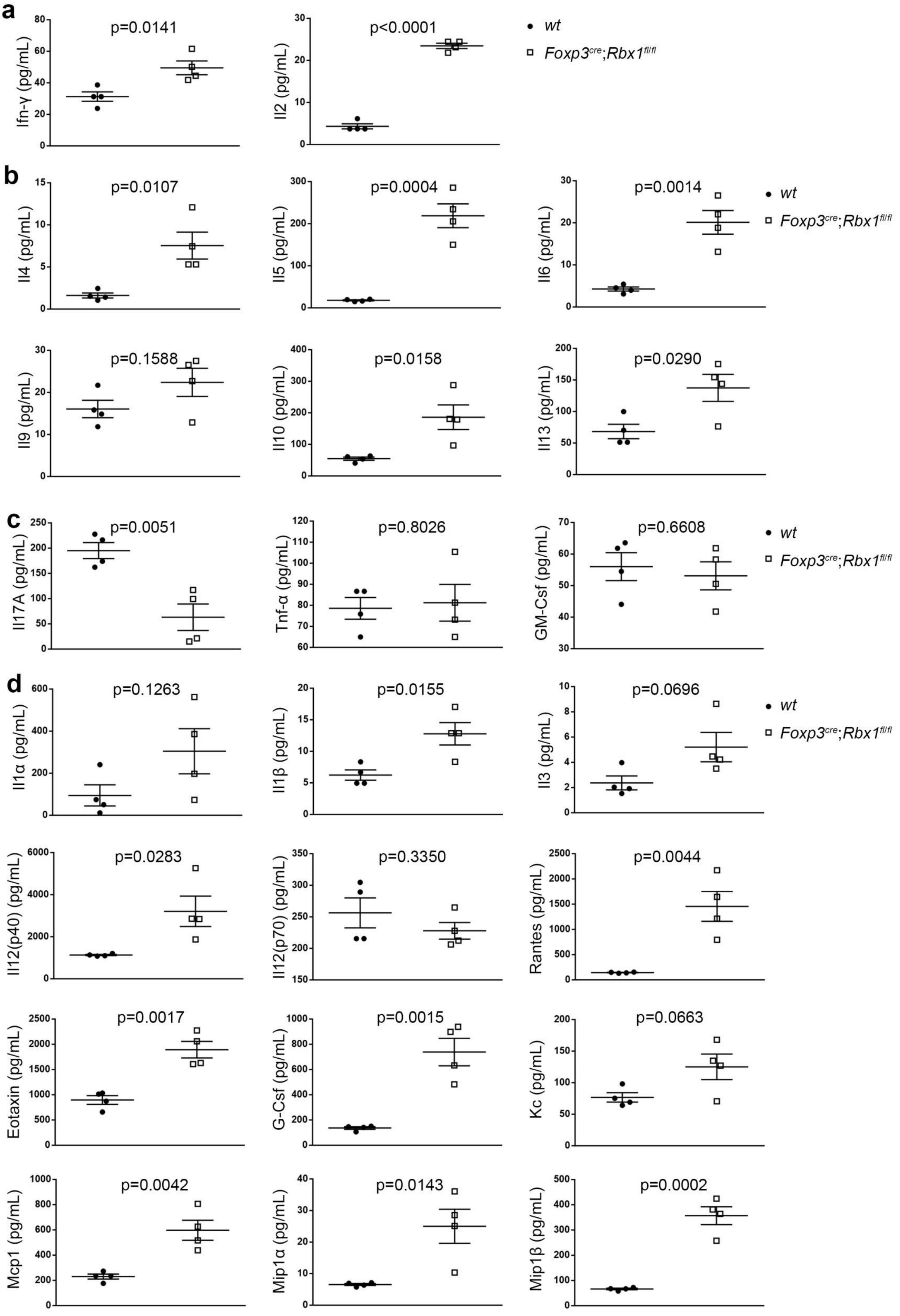
a
p=0.0141
Ifn-γ (pg/mL)
p<0.0001
Il2 (pg/mL)
wt
Foxp3cre;Rbx1fl/fl
b
p=0.0107
Il4 (pg/mL)
p=0.0004
Il5 (pg/mL)
p=0.0014
Il6 (pg/mL)
wt
Foxp3cre;Rbx1fl/fl
p=0.1588
Il9 (pg/mL)
p=0.0158
Il10 (pg/mL)
p=0.0290
Il13 (pg/mL)
c
p=0.0051
Il17A (pg/mL)
p=0.8026
Tnf-α (pg/mL)
p=0.6608
GM-Csf (pg/mL)
wt
Foxp3cre;Rbx1fl/fl
d
p=0.1263
Il1α (pg/mL)
p=0.0155
Il1β (pg/mL)
p=0.0696
Il3 (pg/mL)
wt
Foxp3cre;Rbx1fl/fl
p=0.0283
Il12(p40) (pg/mL)
p=0.3350
Il12(p70) (pg/mL)
p=0.0044
Rantes (pg/mL)
p=0.0017
Eotaxin (pg/mL)
p=0.0015
G-Csf (pg/mL)
p=0.0663
Kc (pg/mL)
p=0.0042
Mcp1 (pg/mL)
p=0.0143
Mip1α (pg/mL)
p=0.0002
Mip1β (pg/mL)

**Extended Data Figure 4. Quantification of serum cytokines in *wt* and *Foxp3^cre^*;*Rbx1^fl/fl^* mice (p16-19, *n* =4)**

a. $T_H1$ cytokines.

b. $T_H2$ cytokines.

c. $T_H17$ cytokines.

d. Other cytokines.

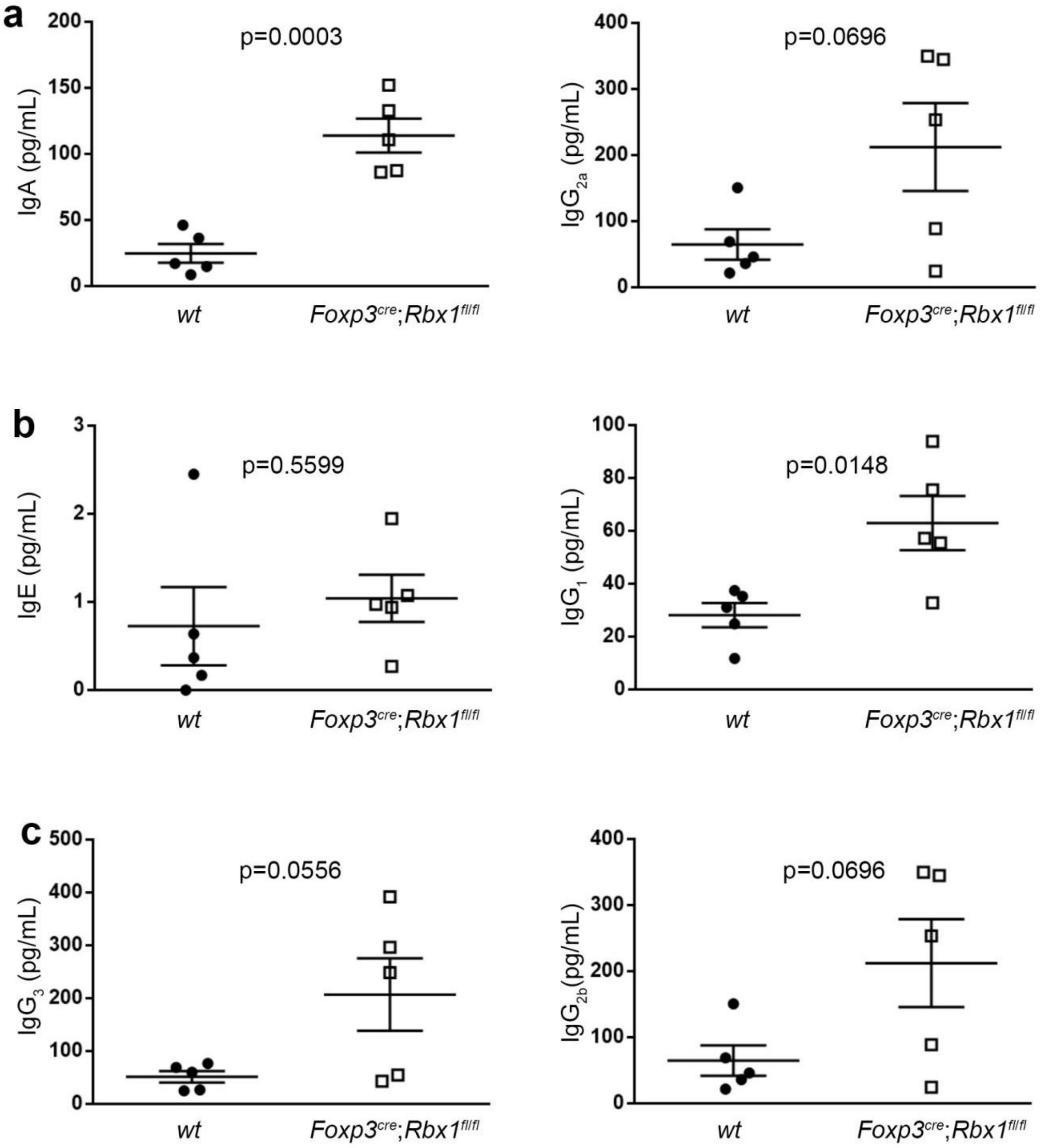


**Extended Data Figure 5. Quantification of serum immunoglobulin subclasses in *wt* and *Foxp3$^{cre}$*;*Rbx1$^{fl/fl}$* mice (p20-21, *n* =5)**

a. $T_H1$ antibodies.

b. $T_H2$ antibodies.

c. $T_H17$ antibodies.

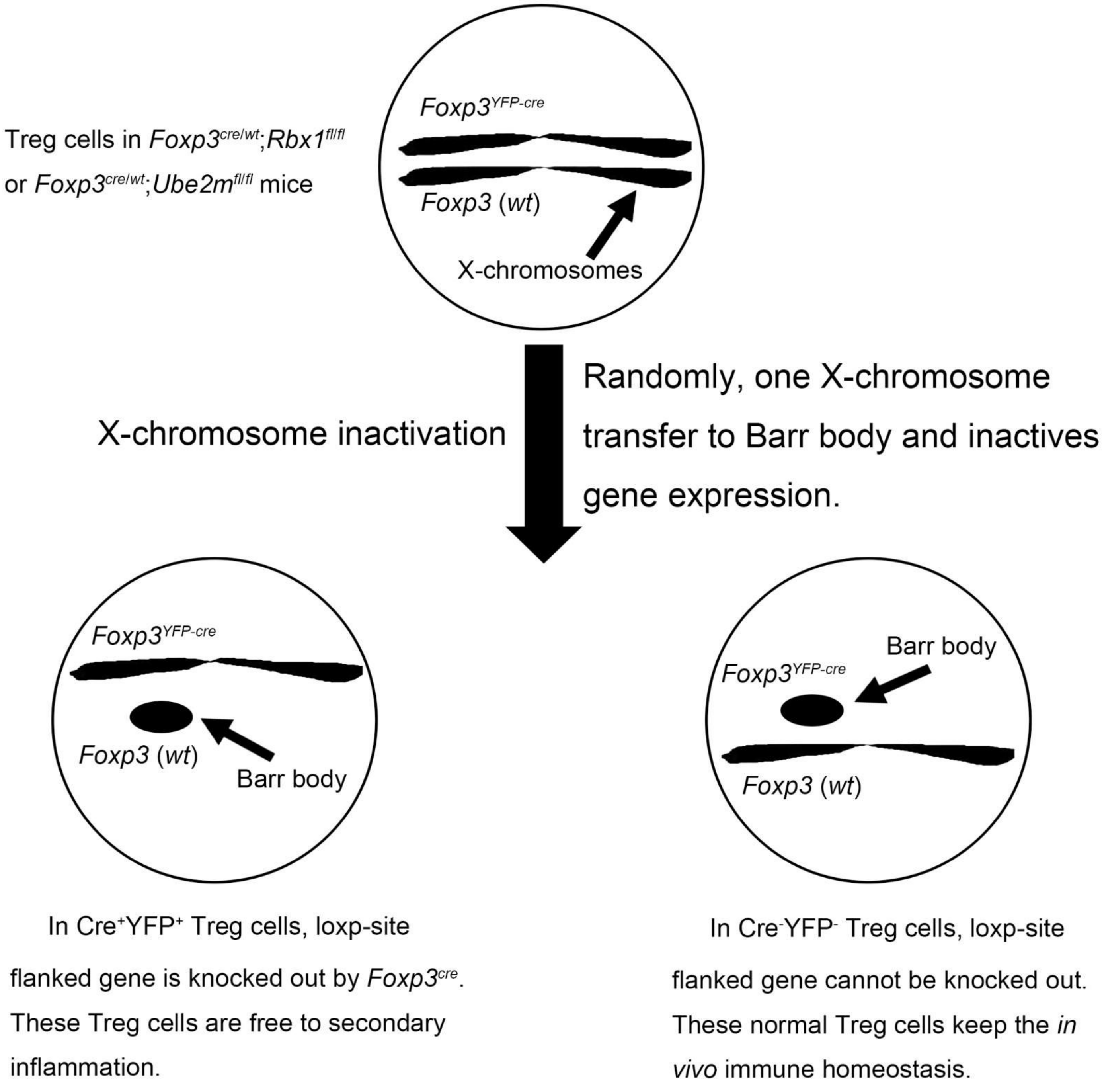


**Extended Data Figure 6. Schematic demonstration how random X-chromosome inactivation protects *Foxp3$^{cre/wt}$*;*Rbx1$^{fl/fl}$* (or Foxp3$^{cre/wt}$;*Ube2m$^{fl/fl}$*) mice from inflammation**

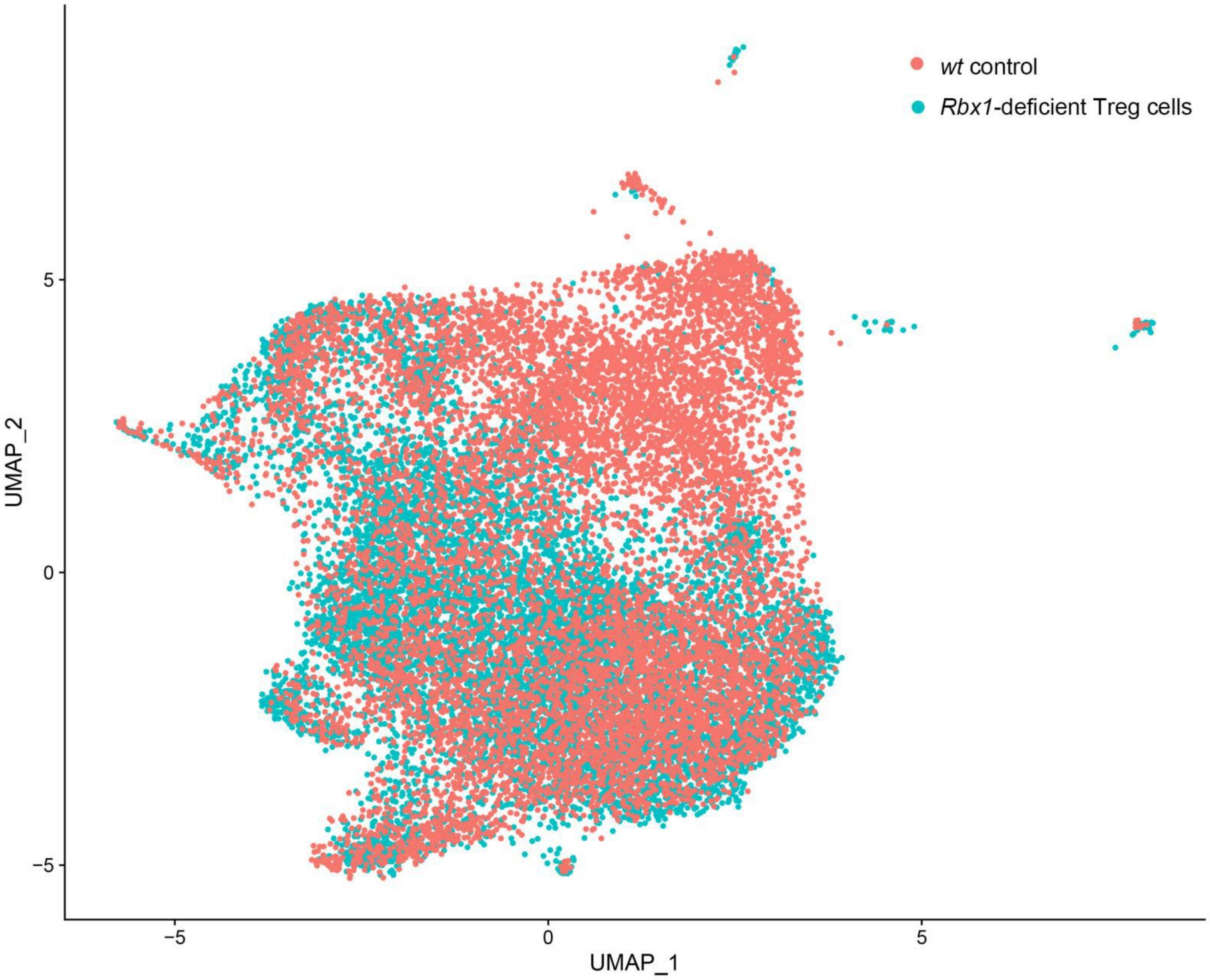


**Extended Data Figure 7. The distribution of *wt* and *Rbx1*-deficient Treg cells in UMAP visualization**

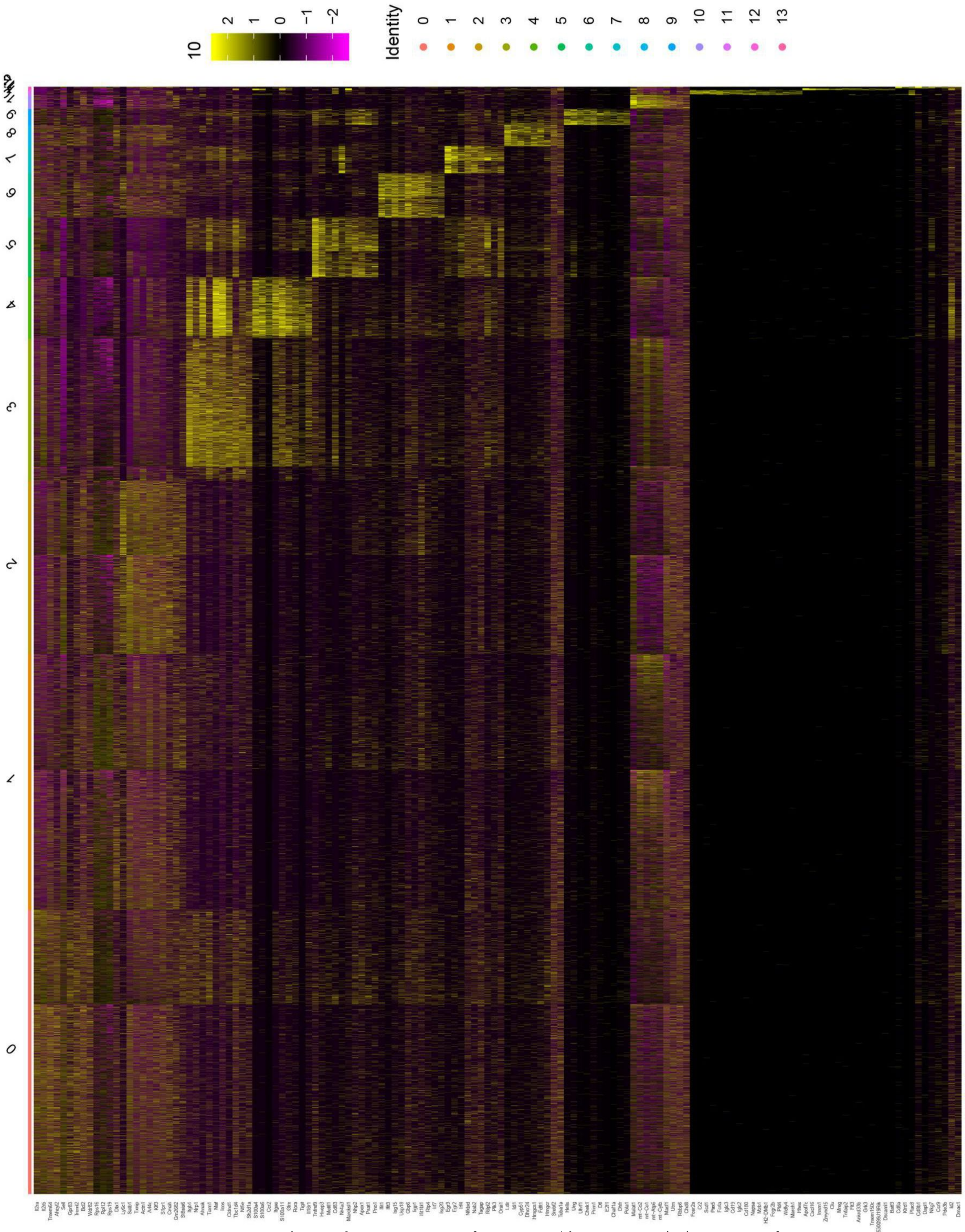


**Extended Data Figure 8. Heat map of the top 10 characteristic genes of each sub-population in *wt* and *Rbx1*-deficient Treg cells**

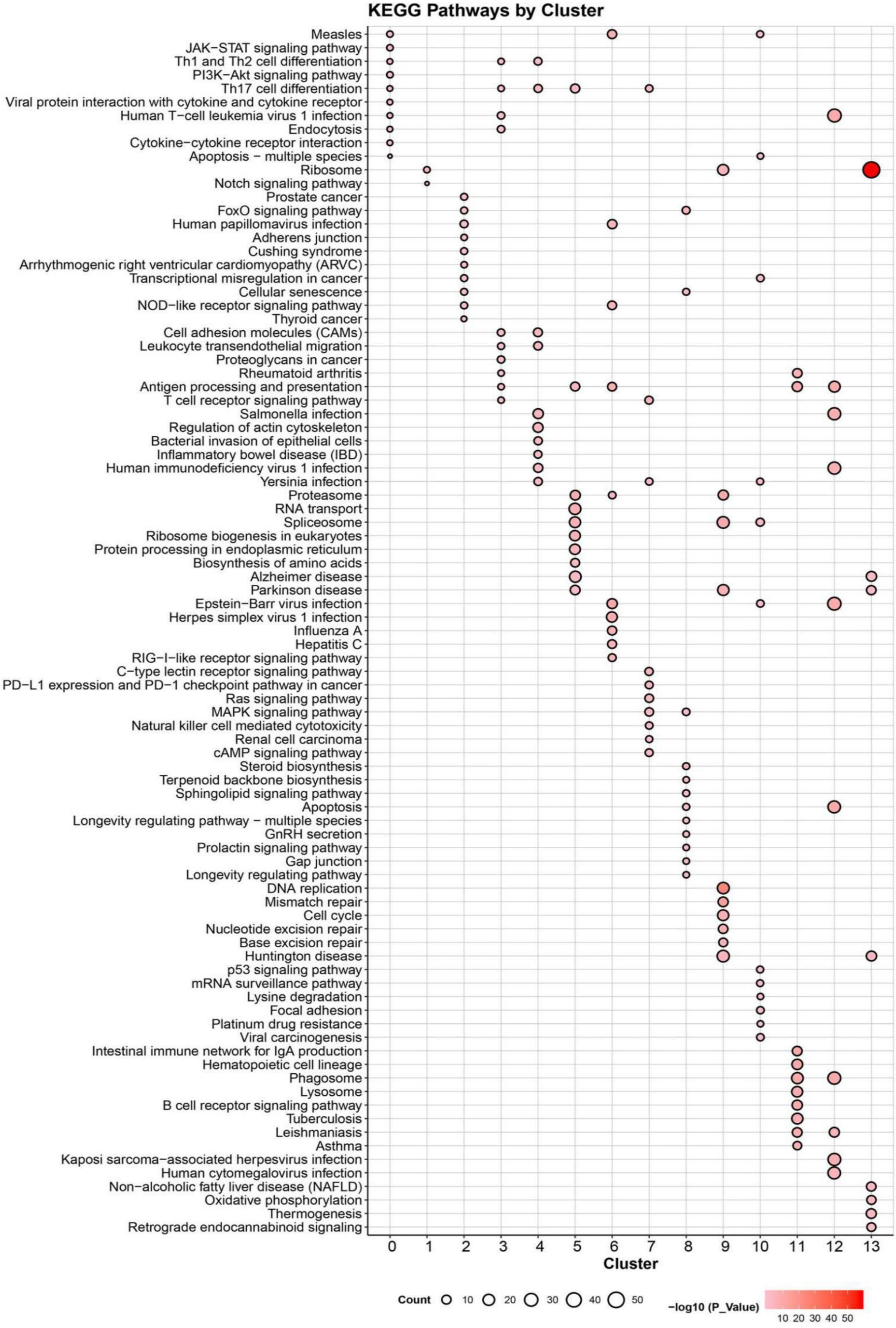


**Extended Data Figure 9. KEGG pathway enrichment of each cluster derived from scRNA-seq analysis of *wt* and *Rbx1*-deficient Treg cells.**

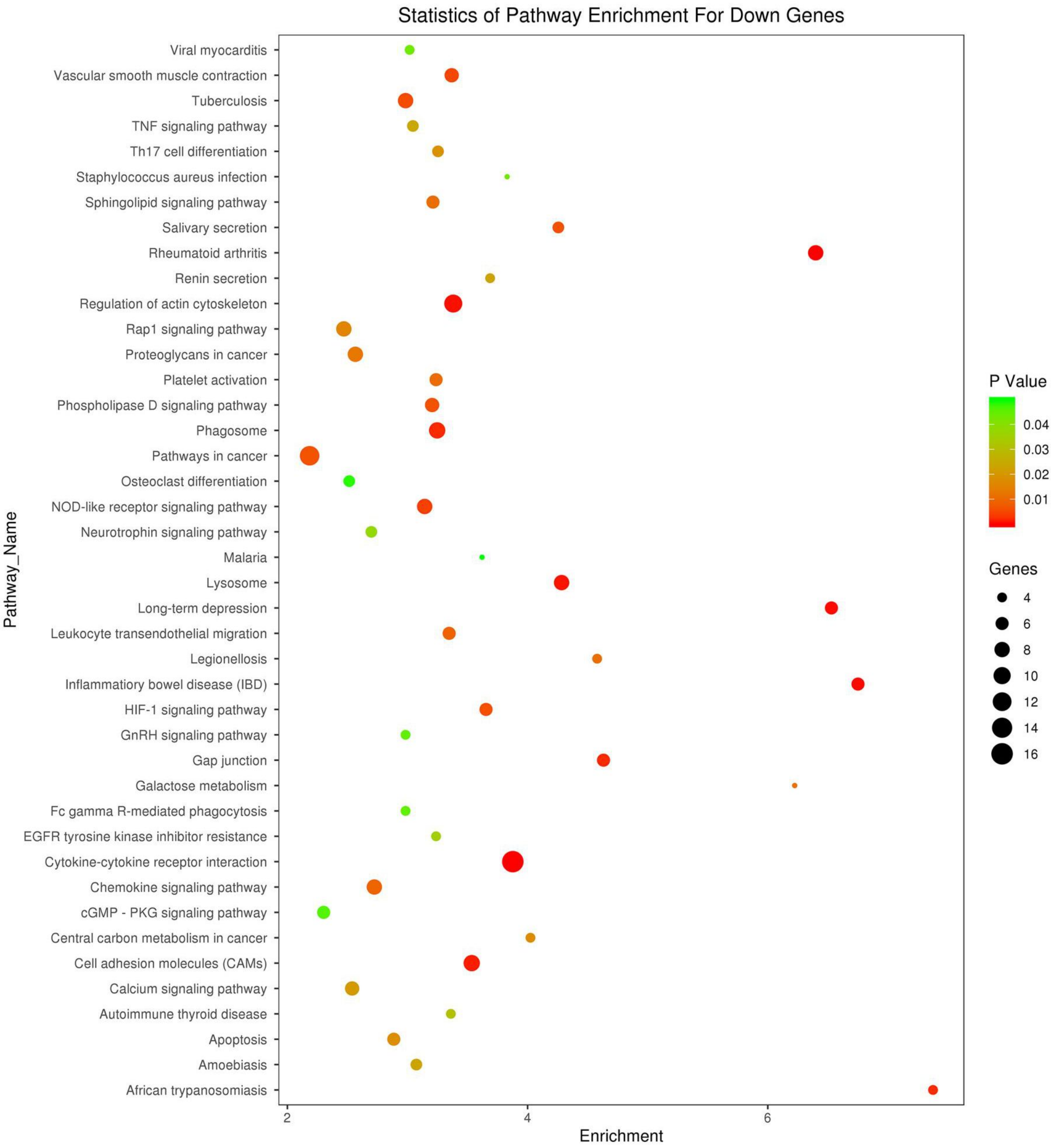


**Extended Data Figure 10. Downregulated pathways in CD4$^{+}$YFP$^{+}$ Treg cells from *Foxp3$^{cre/wt}$* and *Foxp3$^{cre/wt}$;Rbx1$^{fl/fl}$* mice (8-10 weeks old), determined by transcriptional profiling.**

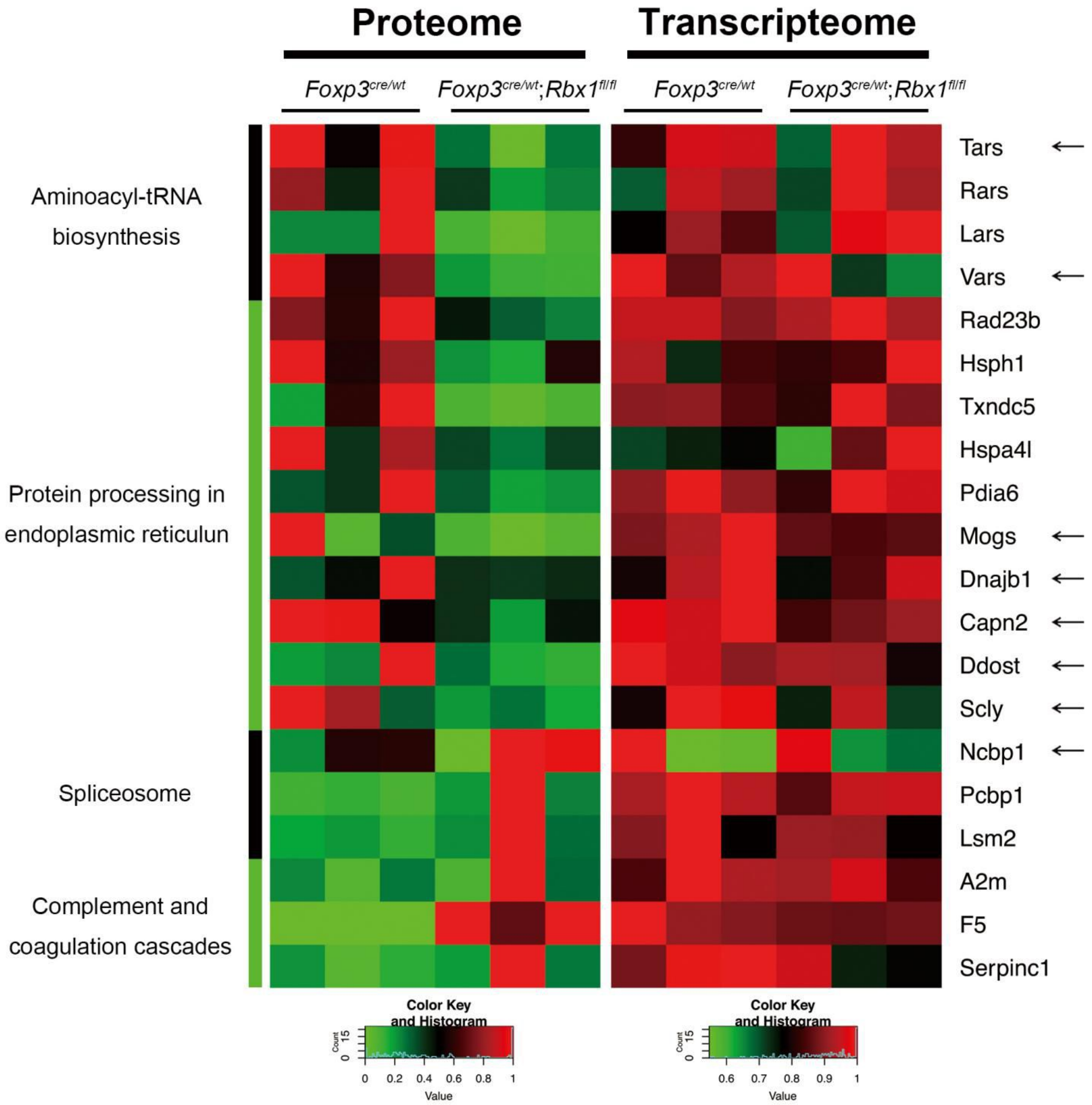


**Extended Data Figure 11. Comparison of the levels of proteins vs. mRNAs in genes associated with indicated pathways in CD4$^{+}$YFP$^{+}$ Treg cells from *Foxp3$^{cre/wt}$* and *Foxp3$^{cre/wt}$;Rbx1$^{fl/fl}$* mice (8-10 weeks old).** The mRNAs with greater than 1.3-fold changes, which are consistent with protein level changes, were marked by the arrows.

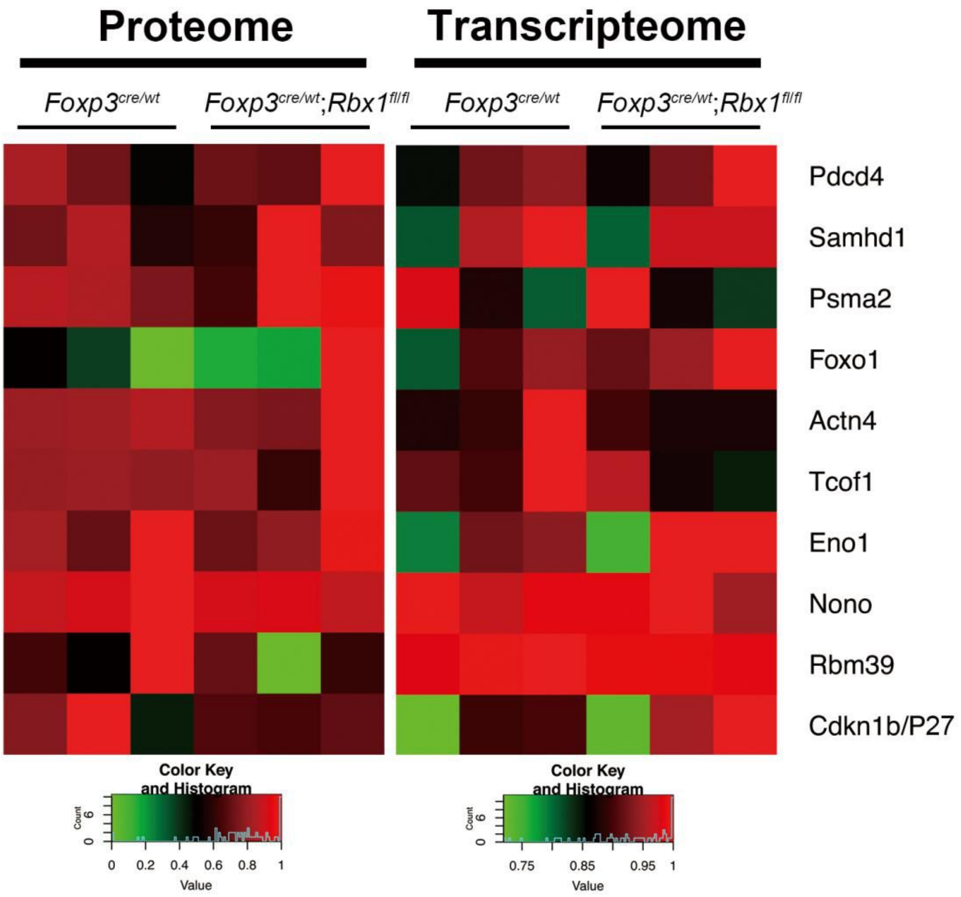


**Extended Data Figure 12. Comparison of the levels of proteins vs. mRNAs of indicated genes in CD4$^{+}$YFP$^{+}$ Treg cells from *Foxp3$^{cre/wt}$* and *Foxp3$^{cre/wt}$;Rbx1$^{fl/fl}$* mice (8-10 weeks old).** Some common substrates of Rbx1 do not accumulated in Treg cells.

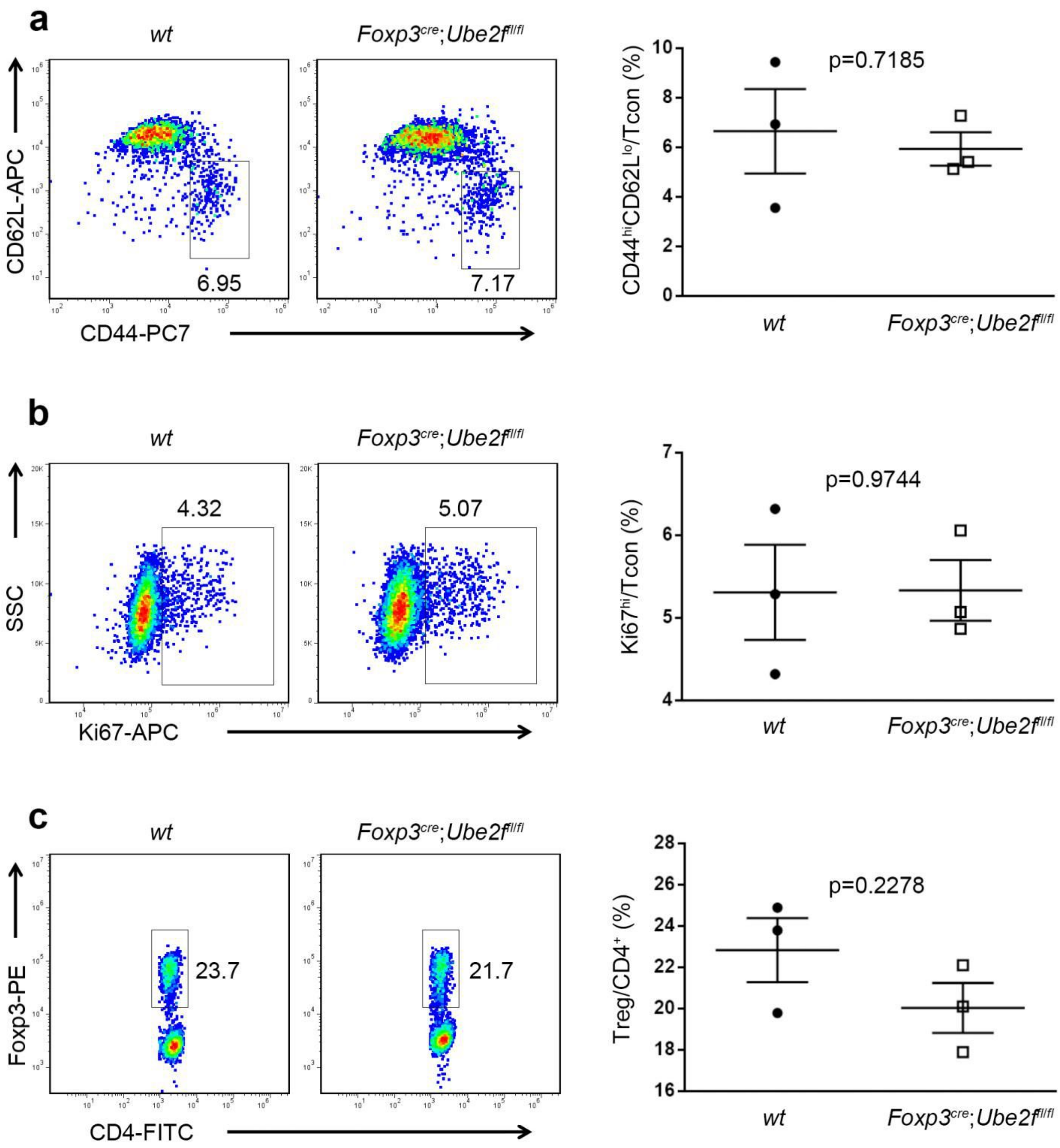


**Extended Data Figure 13. Deficiency of Ube2f does not obviously impair Treg cell fitness at the steady status**

a. Expression of CD44 and CD62L in Tcon cells from peripheral lymph nodes of *wt* and *Foxp3*$^{cre}$;*Ube2f*$^{fl/fl}$ mice (16 weeks old, *n*=3).

b. Expression of Ki67 in Tcon cells in peripheral lymph nodes from *wt* and *Foxp3*$^{cre}$;*Ube2f*$^{fl/fl}$ mice (16 weeks old, *n*=3).

c. The proportion of Treg cells among CD4$^{+}$-T cells in peripheral lymph nodes from *wt* and *Foxp3*$^{cre}$;*Ube2f*$^{fl/fl}$ mice (16 weeks old, *n*=3).

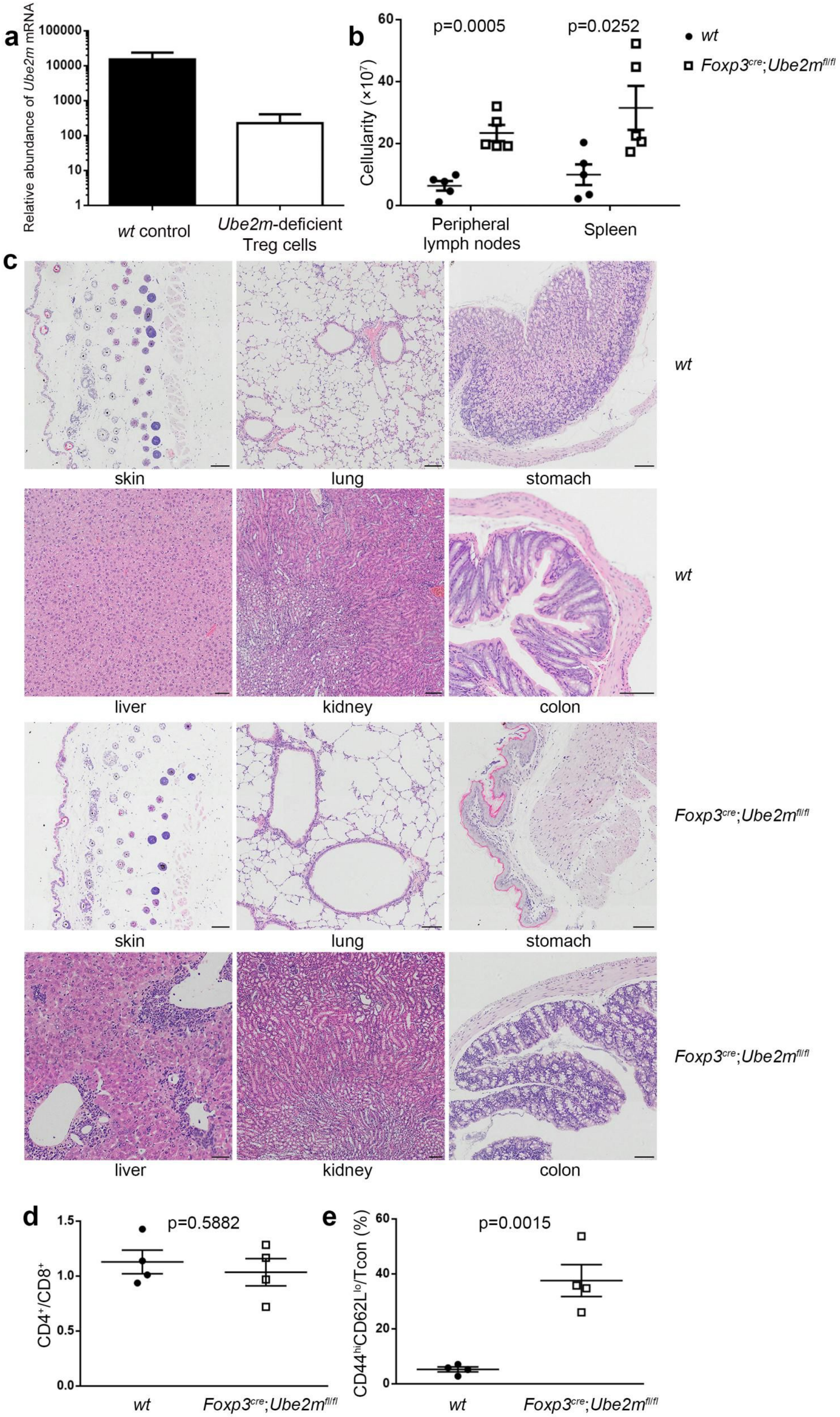

a
Relative abundance of Ube2m mRNA
100000
10000
1000
100
10
1
wt control
Ube2m-deficient Treg cells
b
Cellularity (×10^7)
60
40
20
0
p=0.0005
p=0.0252
Peripheral lymph nodes
Spleen
wt
Foxp3cre;Ube2mfl/fl
c
skin
lung
stomach
wt
liver
kidney
colon
wt
skin
lung
stomach
Foxp3cre;Ube2mfl/fl
liver
kidney
colon
Foxp3cre;Ube2mfl/fl
d
CD4+/CD8+
1.5
1.0
0.5
0.0
p=0.5882
wt
Foxp3cre;Ube2mfl/fl
e
CD44hiCD62Llo/Tcon (%)
60
40
20
0
p=0.0015
wt
Foxp3cre;Ube2mfl/fl

**Extended Data Figure 14. *Ube2m* deletion in Treg cells leads to inflammatory disorders in mice**

a. Relative abundance of *Ube2m* mRNA in *wt* and Ube2m-deficient Treg cells, revealed by transcriptome profiling.

b. Total cell numbers in peripheral lymph nodes and spleen from *wt* and $Foxp3^{cre}$;$Ube2m^{fl/fl}$ mice (16 weeks old, *n*=5).

c. H&E staining of skin, lung, stomach, liver, kidney, colon from *wt* and $Foxp3^{cre}$;$Ube2m^{fl/fl}$ mice (16 weeks old, scale bar = 50µm in liver, and 100µm in other organs).

d. $CD4^+/CD8^+$ ratios in peripheral lymph nodes from *wt* and $Foxp3^{cre}$;$Ube2m^{fl/fl}$ mice (16 weeks old, $n = 4$).

e. $CD44^{hi}CD62L^{lo}$/Tcon ratios in peripheral lymph nodes from *wt* and $Foxp3^{cre}$;$Ube2m^{fl/fl}$ mice (16 weeks old, $n = 4$).

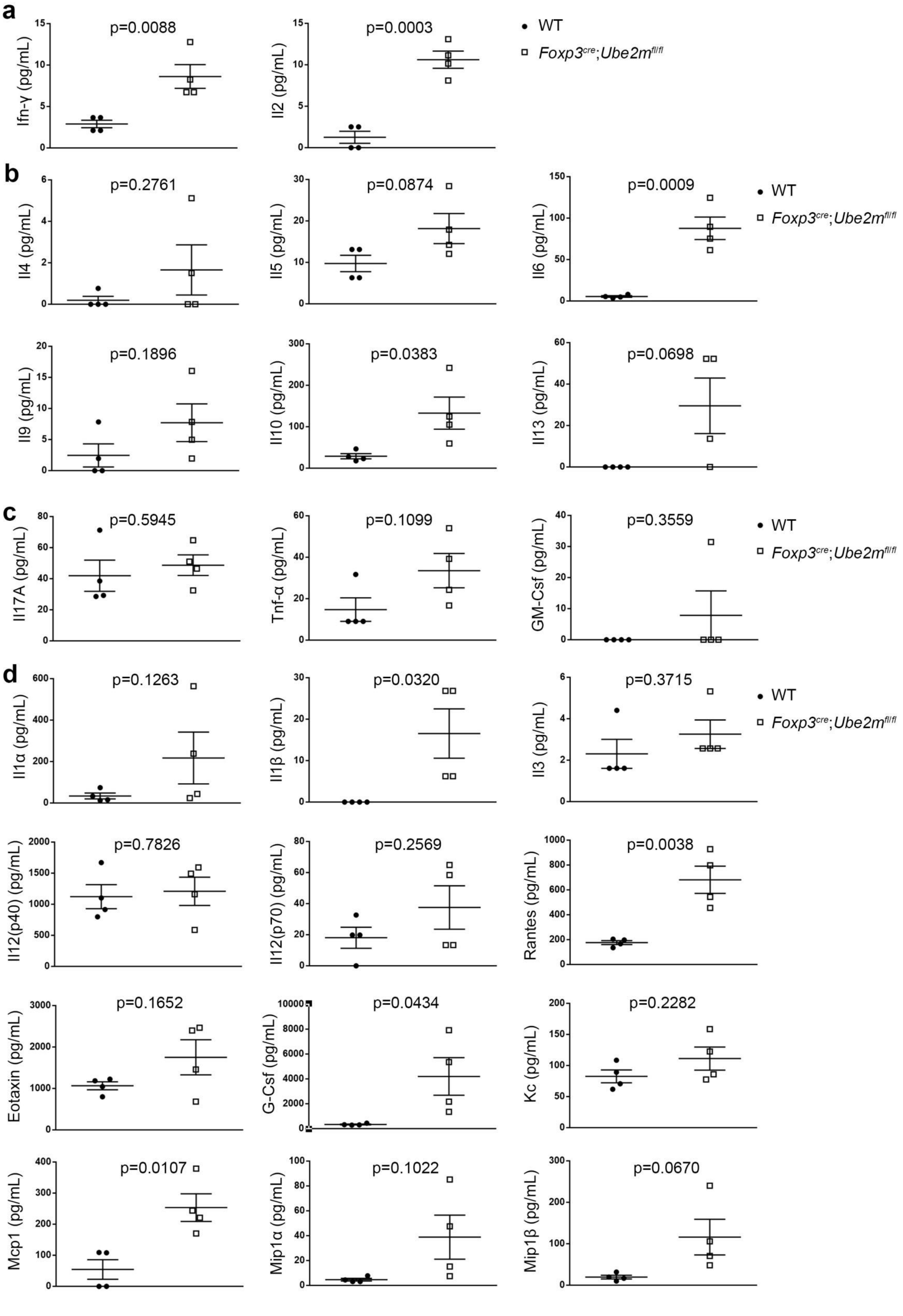
a
b
c
d
WT
Foxp3cre;Ube2mfl/fl
p=0.0088
Ifn-γ (pg/mL)
p=0.0003
Il2 (pg/mL)
p=0.2761
Il4 (pg/mL)
p=0.0874
Il5 (pg/mL)
p=0.0009
Il6 (pg/mL)
p=0.1896
Il9 (pg/mL)
p=0.0383
Il10 (pg/mL)
p=0.0698
Il13 (pg/mL)
p=0.5945
Il17A (pg/mL)
p=0.1099
Tnf-α (pg/mL)
p=0.3559
GM-Csf (pg/mL)
p=0.1263
Il1α (pg/mL)
p=0.0320
Il1β (pg/mL)
p=0.3715
Il3 (pg/mL)
p=0.7826
Il12(p40) (pg/mL)
p=0.2569
Il12(p70) (pg/mL)
p=0.0038
Rantes (pg/mL)
p=0.1652
Eotaxin (pg/mL)
p=0.0434
G-Csf (pg/mL)
p=0.2282
Kc (pg/mL)
p=0.0107
Mcp1 (pg/mL)
p=0.1022
Mip1α (pg/mL)
p=0.0670
Mip1β (pg/mL)

**Extended Data Figure 15. Quantification of serum cytokines in *wt* and *Foxp3$^{cre}$*;*Ube2m$^{fl/fl}$* mice (16 weeks old, *n* =4)**

a. $T_H1$ cytokines.

b. $T_H2$ cytokines.

c. $T_H17$ cytokines.

d. Other cytokines.

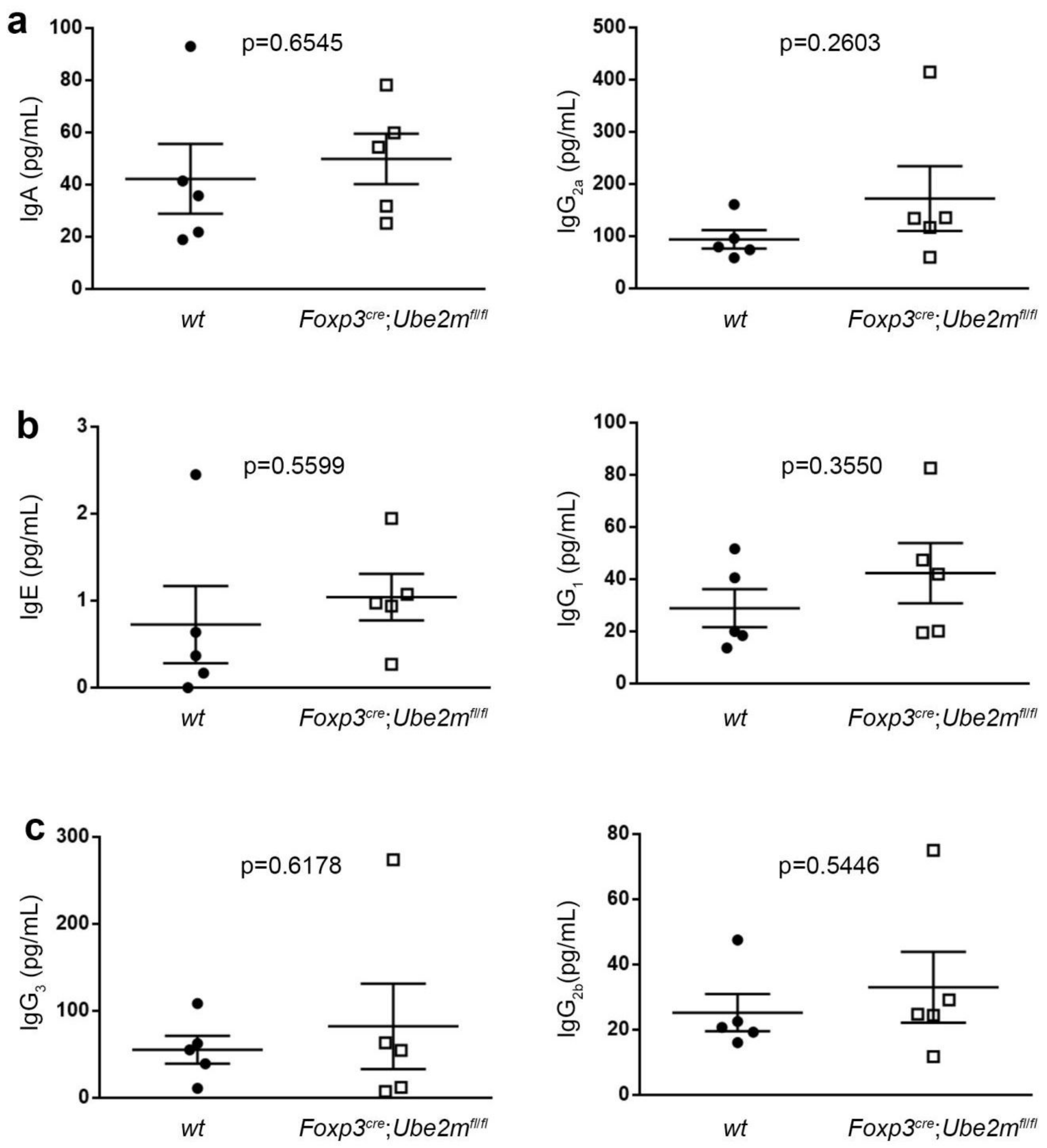


**Extended Data Figure 16. Quantification of serum immunoglobulin subclasses in *wt* and *Foxp3*$^{cre}$;*Ube2m*$^{fl/fl}$ mice (16 weeks old, *n* =5)**

a. $T_H1$ antibodies.

b. $T_H2$ antibodies.

c. $T_H17$ antibodies.

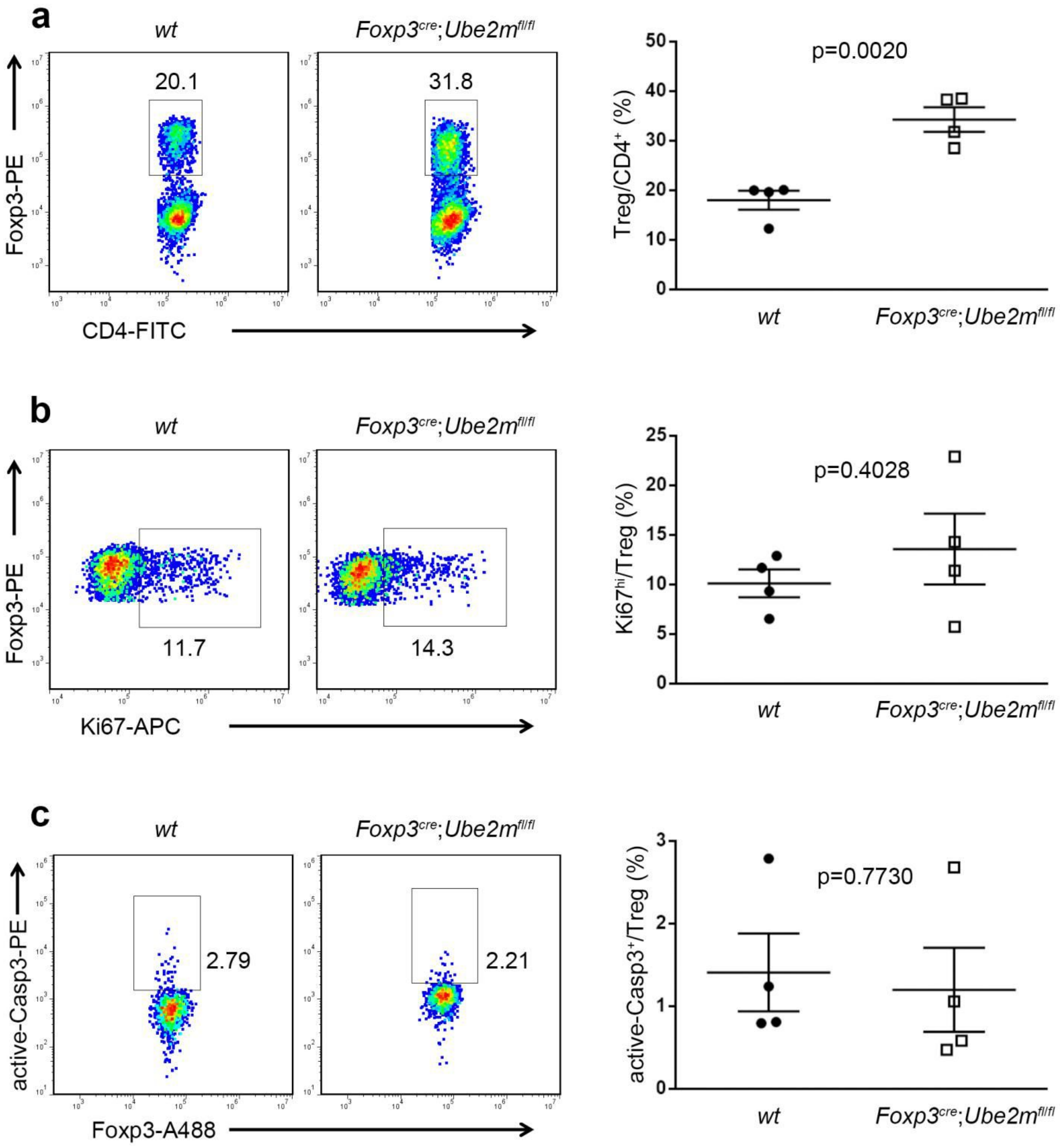


**Extended Data Figure 17. Ube2m-deficiency does not impair Treg cell ratio, proliferation and apoptosis**

a. Treg/$CD4^+$ ratios in peripheral lymph nodes from *wt* and *Foxp3$^{cre}$*;*Ube2m$^{fl/fl}$* mice (16weeks old, *n* =4).

b. Expression of Ki67 in Treg cells in peripheral lymph nodes from *wt* and *Foxp3$^{cre}$*;*Ube2m$^{fl/fl}$* mice (16weeks old, *n* =4).

c. Active Casp3 in Treg cells in peripheral lymph nodes from *wt* and *Foxp3$^{cre}$*;*Ube2m$^{fl/fl}$* mice (16weeks old, *n* =4).

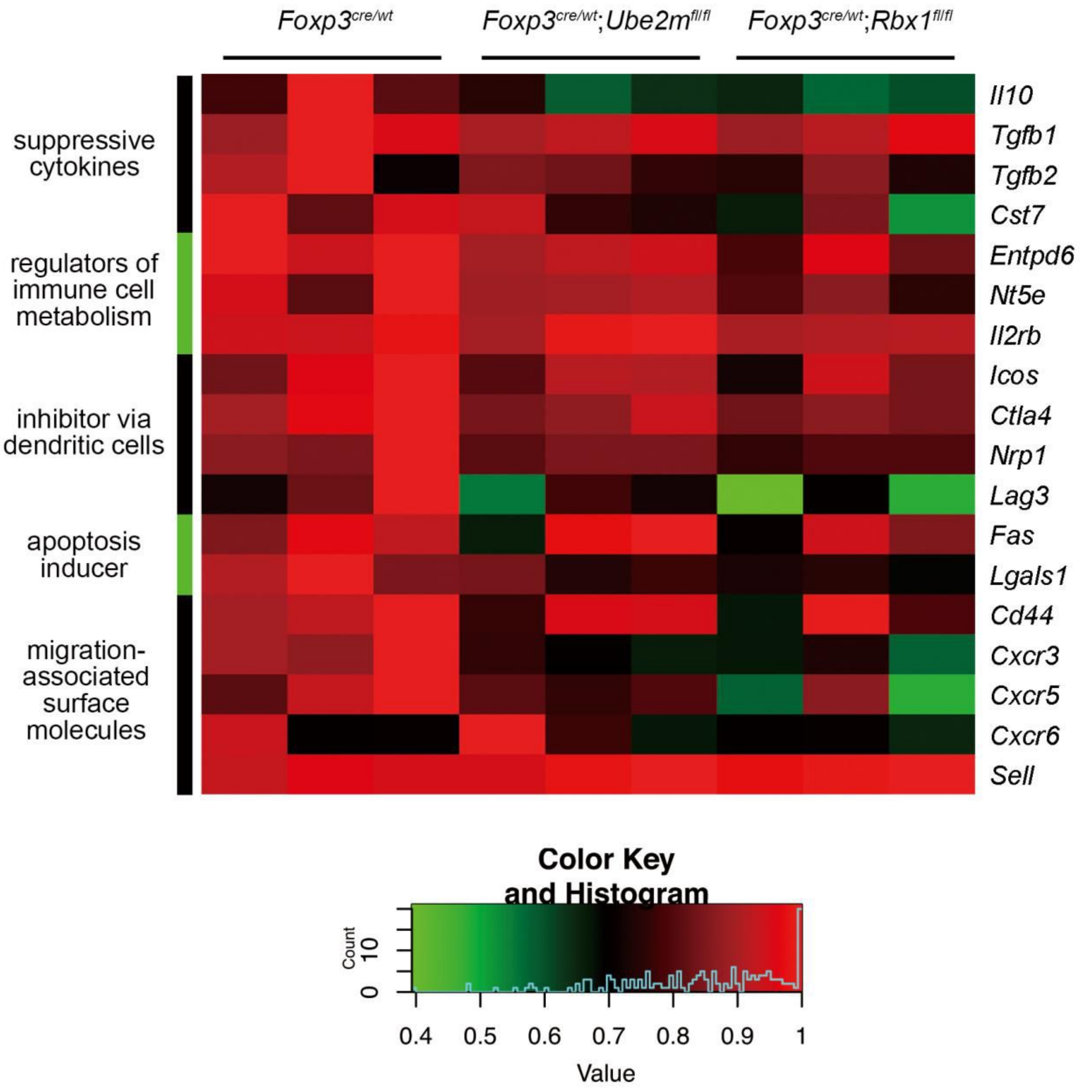


**Extended Data Figure 18. The expression levels of indicated Treg cell function-related genes in CD4$^{+}$YFP$^{+}$ Treg cells derived from *Foxp3$^{cre/wt}$*, *Foxp3$^{cre/wt}$;Ube2m$^{fl/fl}$* and *Foxp3$^{cre/wt}$;Rbx1$^{fl/fl}$* mice (8-10 weeks old), determined by transcriptional profiling.**

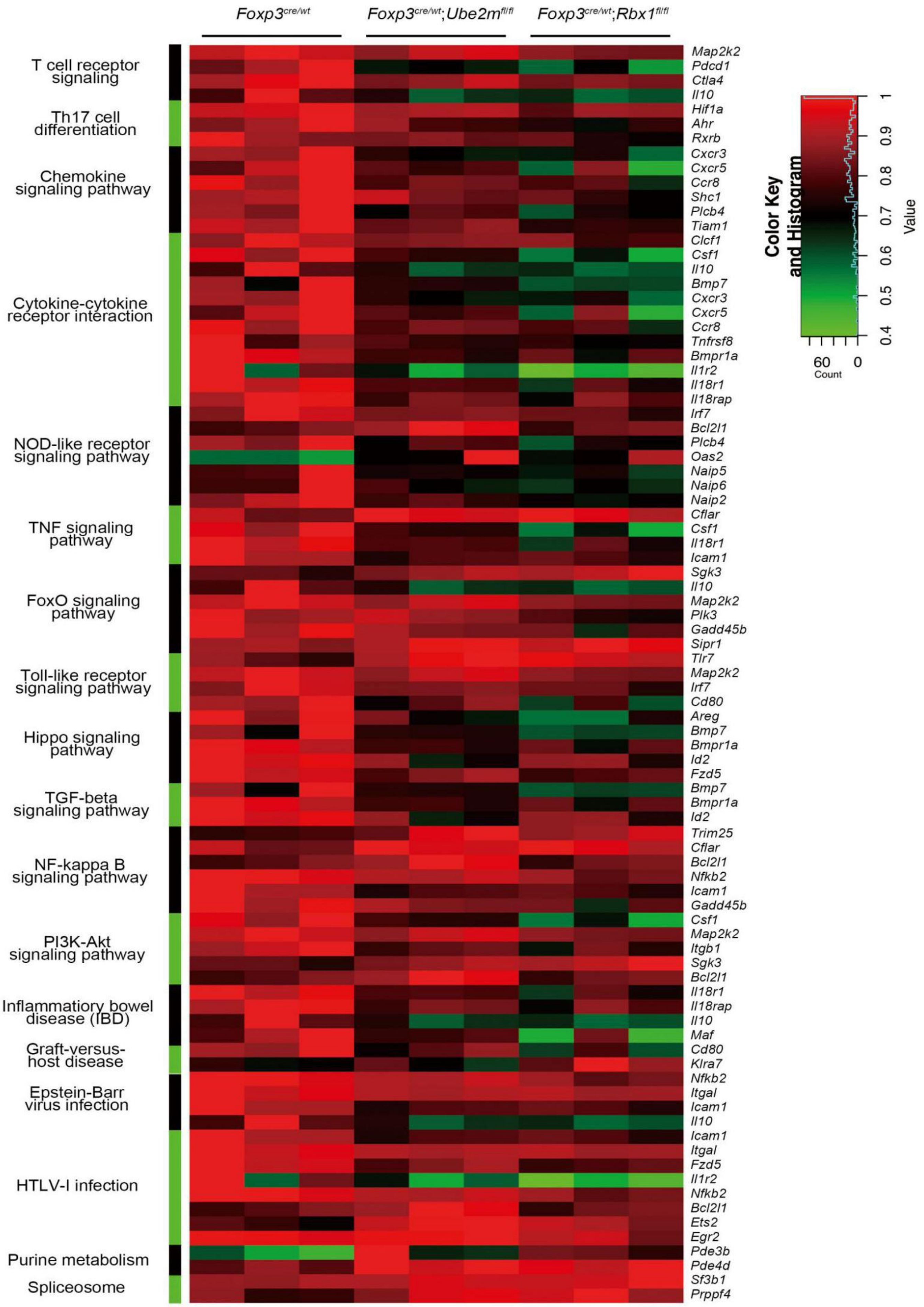


**Extended Data Figure 19. Comparison of expression levels of indicated genes and their associated signaling pathways in $CD4^{+}YFP^{+}$ Treg cells derived from *Foxp3*$^{cre/wt}$, *Foxp3*$^{cre/wt}$;*Ube2m*$^{fl/fl}$ and *Foxp3*$^{cre/wt}$;*Rbx1*$^{fl/fl}$ mice (8-10 weeks old), determined by transcriptional profiling.**

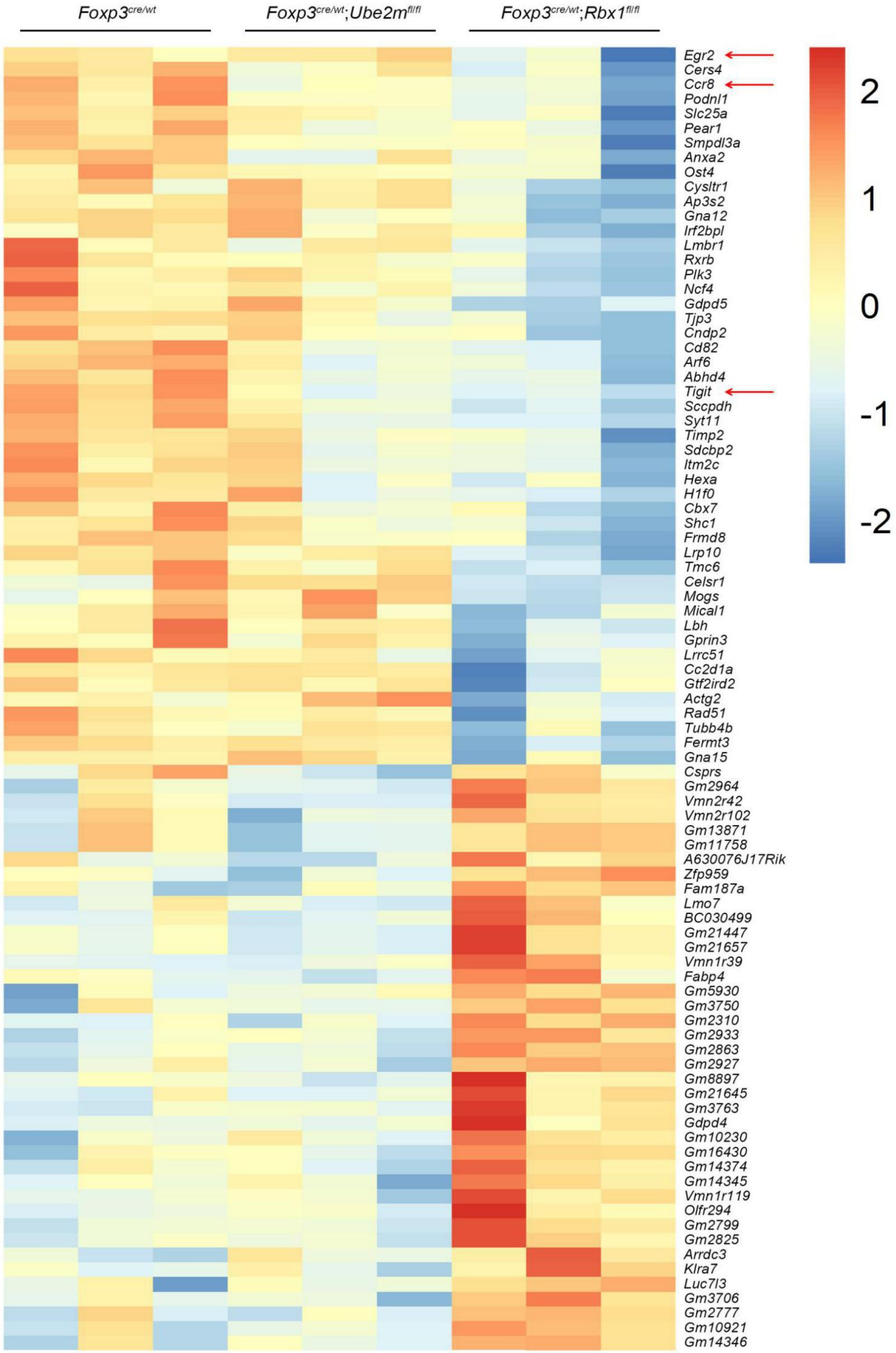


**Extended Data Figure 20. Genes with altered expression selectively in Rbx1-deficient Treg cells.** Top portion: Genes with reduced expression; Bottom panels: Genes with increased expression.

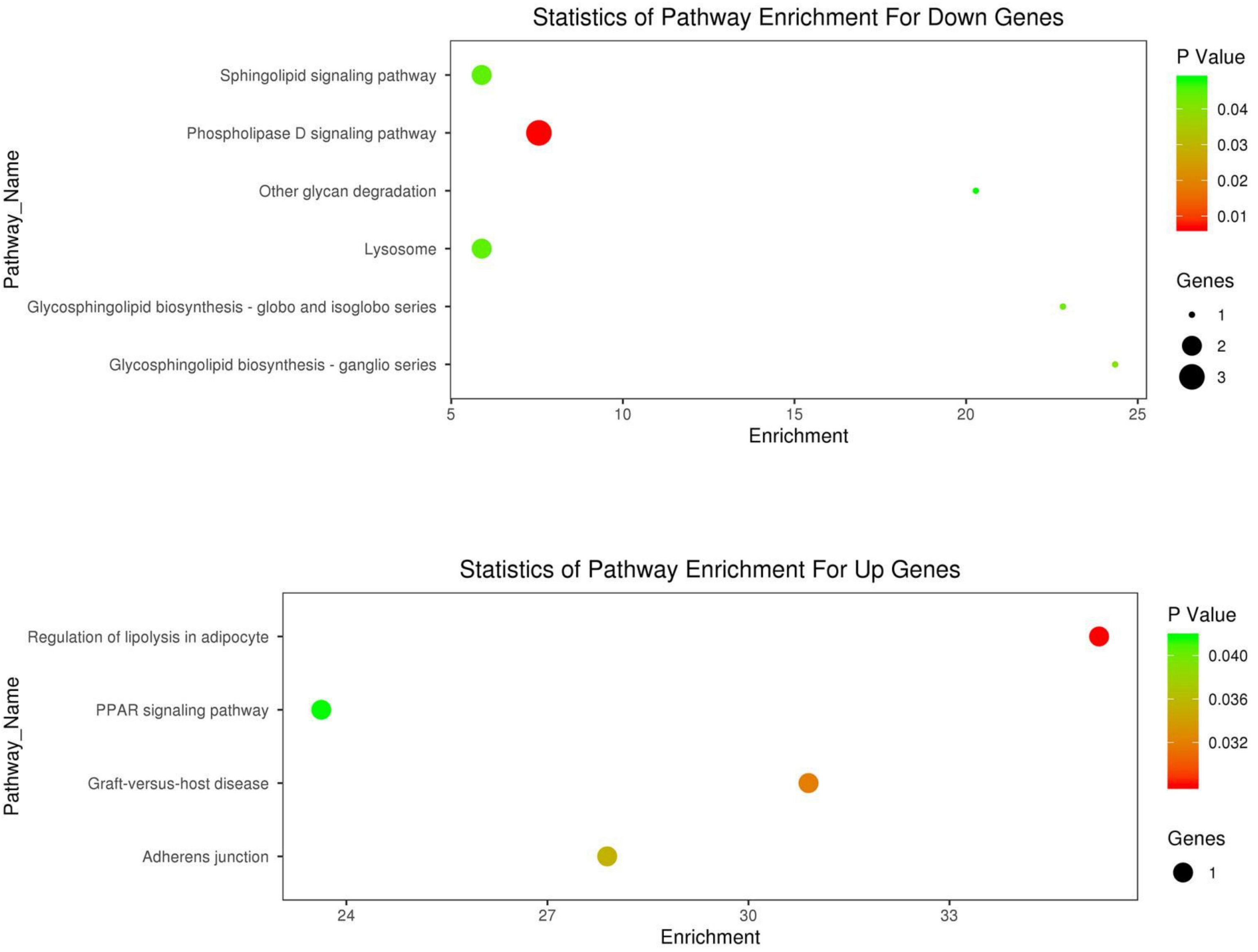


**Extended Data Figure 21. Annotation of KEGG pathways selectively altered in Rbx1-deficient Treg cells.** Top panel: enriched pathways from down-regulated genes; Bottom panel: enriched pathways from up-regulated genes.
